# Multimodal decoding of human liver regeneration

**DOI:** 10.1101/2023.02.24.529873

**Authors:** KP Matchett, JR Wilson-Kanamori, JR Portman, CA Kapourani, F Fercoq, S May, JBG Mackey, M Brice, E Zajdel, M Beltran, EF Sutherland, GC Wilson, SJ Wallace, L Kitto, NT Younger, R Dobie, GC Oniscu, SJ Wigmore, P Ramachandran, CA Vallejos, NO Carragher, KJ Simpson, TJ Kendall, Acute Liver Failure Study Group, JA Rule, WM Lee, M Hoare, CJ Weston, JC Marioni, ST Teichmann, TG Bird, LM Carlin, NC Henderson

**Author notes:** Address correspondence to: Neil Henderson, Centre for Inflammation Research, The Queen’s Medical Research Institute, University of Edinburgh, Edinburgh, UK. These authors contributed equally (JR Wilson-Kanamori, JR Portman, CA Kapourani).

## Abstract

The liver has a unique ability to regenerate^1,2^, however in the setting of acute liver failure (ALF) this regenerative capacity is often overwhelmed and emergency liver transplantation is the only curative option^3-5^. To advance our understanding of human liver regeneration and to inform design of pro-regenerative therapies, we use paired single-nuclei RNA sequencing (snRNA-seq) combined with spatial profiling of healthy and ALF explant human livers to generate the first single-cell, pan-lineage atlas of human liver regeneration. We uncover a novel ANXA2^+^ migratory hepatocyte subpopulation which emerges during human liver regeneration, and a corollary migratory hepatocyte subpopulation in a mouse model of acetaminophen (APAP)-induced liver regeneration. Importantly, interrogation of necrotic wound closure and hepatocyte proliferation across multiple timepoints following APAP-induced liver injury in mice demonstrates that wound closure precedes hepatocyte proliferation. 4-D intravital imaging of APAP-induced mouse liver injury identifies motile hepatocytes at the edge of the necrotic area, enabling collective migration of the hepatocyte sheet to effect wound closure. Depletion of hepatocyte ANXA2 expression reduces HGF-induced human and mouse hepatocyte migration *in vitro*, and abrogates necrotic wound closure following APAP-induced mouse liver injury. Taken together, our work dissects unanticipated aspects of liver regeneration, demonstrating an uncoupling of wound closure and hepatocyte proliferation and uncovering a novel migratory hepatocyte subpopulation which mediates wound closure following liver injury. Therapies designed to promote rapid reconstitution of normal hepatic microarchitecture and reparation of the gut-liver barrier may open up new areas of therapeutic discovery in regenerative medicine.

## Main

Acute liver failure (ALF) is a syndrome of severe liver injury in the absence of chronic liver disease^3-5^. ALF is often unexpected, affecting previously healthy individuals, and has a rapid onset with a frequently fatal outcome (30% mortality)^3^. The major causes of ALF in the UK and the USA are acetaminophen (paracetamol) toxicity, nonA-E hepatitis, ischaemia, drug-induced liver injury, hepatitis B virus, and autoimmunity, with acetaminophen toxicity representing the commonest cause of ALF in the UK (65.4% of cases) and USA (45.7% of cases)^3^. In contrast, viral hepatitis A, B, and E are the main causes of ALF in Asia^6^. In severe cases of ALF, emergency liver transplantation remains the only curative option. Therefore, effective pro-regenerative therapies, designed to harness and augment the inherent regenerative and reparative capacity of the liver, are urgently required.

Hepatocytes, the major epithelial component of the liver accounting for ∼80% of its mass, perform a vast array of vital metabolic and synthetic functions and are therefore fundamental in the maintenance of normal liver function^7,8^. Hepatocyte replenishment can occur via conversion of cholangiocytes during severe liver injury^9,10^, however recent studies have demonstrated that hepatocytes are primarily maintained by the proliferation of pre-existing hepatocytes during liver homeostasis and regeneration^11-15^. Despite major advances in our understanding of the modes of hepatocyte replenishment during liver regeneration, a key question remains regarding how the liver restores normal microarchitecture, and hence gut-liver barrier function, following necro-inflammatory liver injury.

Here, using a cross-species, integrative multimodal approach, we have investigated the cellular and molecular mechanisms regulating liver regeneration. Our data define: (1) the first single-cell, pan-lineage atlas of human liver regeneration; (2) a novel ANXA2^+^ migratory hepatocyte subpopulation which emerges during human liver regeneration; (3) a corollary migratory hepatocyte subpopulation in APAP-induced mouse liver injury; (4) that wound closure precedes hepatocyte proliferation during APAP-induced mouse liver injury; (5) motile hepatocytes (assessed using 4-D intravital imaging) at the edge of the necrotic area of APAP-induced mouse liver injury enable collective migration of the hepatocyte sheet to effect wound closure; (6) that depletion of hepatocyte ANXA2 expression reduces hepatocyte growth factor (HGF)-induced human and mouse hepatocyte migration *in vitro*, and abrogates necrotic wound closure following APAP-induced mouse liver injury.

Our work dissects unanticipated aspects of liver regeneration, demonstrating an uncoupling of wound closure and hepatocyte proliferation and uncovering a novel migratory hepatocyte subpopulation which mediates wound closure following liver injury.

## Results

### Deconstructing human liver regeneration

Initially, we screened human liver explant samples from patients transplanted for multiple aetiologies of acute liver failure (acetaminophen-induced and nonA-E hepatitis) and chronic liver disease (non-alcoholic fatty liver disease, alcohol-induced, primary biliary cholangitis and primary sclerosing cholangitis) to investigate which human liver diseases exhibit a substantial hepatocyte proliferative response. Acetaminophen-induced acute liver failure (APAP-ALF) and nonA-E hepatitis (NAE-ALF) demonstrated markedly increased hepatocyte proliferation compared to uninjured human liver and all chronic human liver diseases examined (Fig. 1a). We therefore focussed on APAP-ALF and NAE-ALF in this study of human liver regeneration.

**Figure 1:**
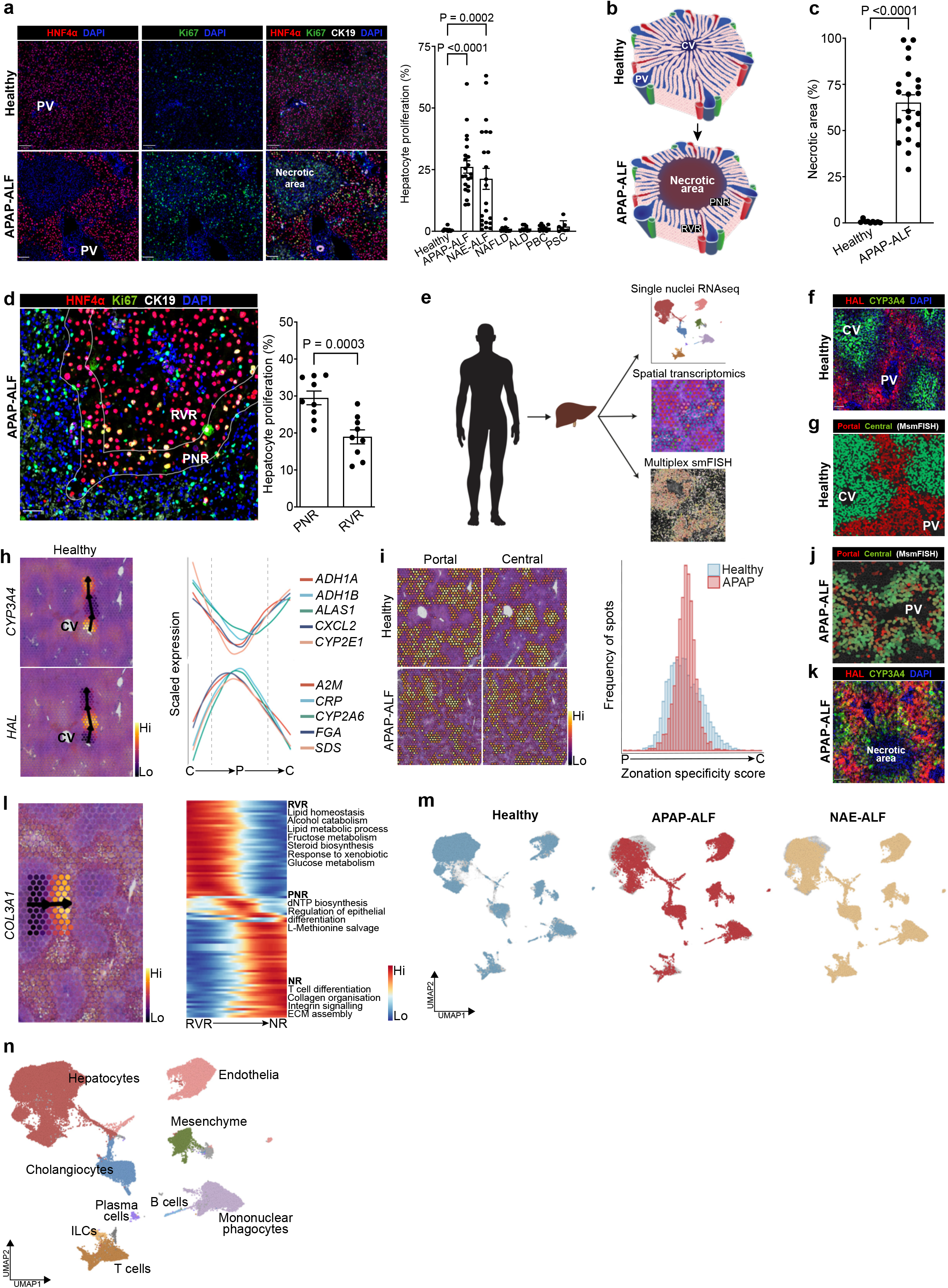
Deconstructing human liver regeneration. **a**, Representative immunofluorescence images of HNF4α (hepatocytes, red), Ki67 (green), CK19 (cholangiocytes, white) and DAPI (nuclear stain, blue) in human healthy and APAP-ALF liver tissue (left). Scale bar 100µm. Hepatocyte proliferation in human healthy and diseased explant livers across multiple aetiologies (healthy n=9, APAP-ALF n=22, NAE-ALF n=22, NAFLD (non-alcoholic fatty liver disease) n=10, ALD (alcohol-induced liver disease) n=9, PBC (primary biliary cholangitis) n=10, PSC (primary sclerosing cholangitis (n=7) (right). One-way ANOVA, *F*=11.46, df=6,82. Data are mean±SEM. **b**, Schematic of the healthy liver lobule. CV, central vein; PV, portal vein (top). Acetaminophen poisoning, if left untreated, can result in massive, confluent necrosis of hepatocytes in the peri-central vein region of the liver lobule (bottom). PNR, peri-necrotic region; RVR, remnant viable region **c**, Quantification of necrotic area in human healthy (n=9) and APAP-ALF (n=22) liver tissue. Unpaired t-test, t=8.76, df=30. Data are mean±SEM. **d**, Representative immunofluorescence image of HNF4α (hepatocytes, red), Ki67 (green), CK19 (cholangiocytes, white) and DAPI (blue) in human APAP-ALF liver tissue (left). Scale bar 50µm. Hepatocyte proliferation in the peri-necrotic region (PNR) and remnant viable region (RVR) of human APAP-induced ALF liver tissue (n=9, right). Paired t-test, t=6.20, df=8. Data are mean±SEM. **e**, Schematic of human liver explant tissue processing for single nuclei RNAseq, spatial transcriptomics, and multiplex smFISH. **f**, Representative immunofluorescence image of HAL (portal hepatocytes, red), CYP3A4 (central hepatocytes, green) and DAPI (nuclear stain, blue) in healthy human liver tissue. Scale bar 100µm. **g**, Spatial expression (multiplex smFISH) of known hepatocyte zonation gene modules in healthy human liver tissue. Gene modules in SI Table 2. **h**, Representative spatial trajectory analysis to identify differentially expressed gene modules across the healthy human liver lobule. **i**, Spatial expression (ST) of healthy human liver-derived zonal gene modules in healthy and APAP-ALF liver tissue (left). Distribution of zonal specificity score in healthy and APAP-ALF liver tissue (right). **j**, Spatial expression (multiplex smFISH) of known hepatocyte zonation gene modules in human APAP-ALF liver tissue. Gene modules in SI Table 2. **k**, Representative immunofluorescence image of HAL (portal hepatocytes, red), CYP3A4 (central hepatocytes, green) and DAPI (nuclear stain, blue) in human APAP-ALF liver tissue. Scale bar 100µm. **l**, Representative spatial trajectory analysis (left) followed by differential GO terms across the human APAP-ALF liver lobule (right). **m**, UMAP visualisation of 72,262 nuclei from healthy (n=9), APAP-ALF (n=10), and NAE-ALF (n=12) human liver explants. **n**, UMAP of cell lineage inferred using signatures of known lineage markers. ILCs, innate lymphoid cells.

Structurally, the healthy liver is divided into lobules, with each lobule consisting of a central vein surrounded by portal tracts (each containing a portal vein, hepatic artery, and bile duct; Fig. 1b, upper). NAE-ALF, a disease of unknown cause, results in massive necrosis of hepatocytes across the lobule. In contrast, severe acetaminophen poisoning can result in massive, confluent necrosis of hepatocytes (Fig. 1c) extending out from the peri-central vein region of the liver lobule (Fig. 1b, lower). Hepatocyte proliferation in human APAP-ALF was increased in the peri-necrotic region (PNR) compared to the residual viable region (RVR; Fig. 1d).

To deconstruct human liver regeneration we applied a multimodal approach (Fig. 1e) including single nuclei RNA sequencing (snRNA-seq), spatial transcriptomics (ST) and multiplex single molecule fluorescence *in situ* hybridization (multiplex smFISH). As transplantation for ALF is much less frequent than transplantation for chronic liver disease, we sourced biobanked, frozen liver explant tissue samples from multiple liver transplant centres in the UK and the USA (Edinburgh, Birmingham, and Cambridge, UK, and the Acute Liver Failure Study Group, USA).

To investigate potential disruption of zonation in human liver regeneration, we spatially profiled liver samples from healthy and APAP-ALF patients (SI Table 1). As expected, ST recapitulated hepatocyte and myofibroblast topography in APAP-ALF versus healthy controls (Extended Data Fig. 1a). ST also showed increased cell cycling in APAP-ALF versus healthy controls (Extended Data Fig. 1a), consistent with the previously observed topography of hepatocyte proliferation in APAP-ALF using immunofluorescence (IF) staining (Fig. 1a,d). Guided by IF staining (Fig. 1f) and multiplex smFISH (Fig. 1g) of established hepatocyte zonation markers, we drew spatial trajectories in ST using the SPATA framework to identify gene modules differentially expressed across the lobule (Fig. 1h, SI Table 2). Gene ontology (GO) trajectory analysis in SPATA confirmed known peri-central and peri-portal hepatocyte biological processes (Extended Data Fig. 1b, SI Table 4). Applying these zonal gene modules to APAP-ALF samples demonstrated loss of hepatocyte portal-central polarity, and emergence of hepatocytes with both portal and central characteristics (Fig. 1i); this disruption in zonation was confirmed using multiplex smFISH (Fig. 1j, Extended Data Fig. 1c,d) and IF staining (Fig. 1k). Further spatial trajectory analysis revealed mixed portal- and central-associated GO terms in the residual viable region of APAP-ALF samples, distinct from those present in peri-necrotic and necrotic regions (Fig. 1l, SI Table 4). These data demonstrate that remnant human hepatocytes display functional plasticity to compensate for substantial loss of peri-central hepatocytes following APAP-ALF.

To further decode human liver regeneration, we performed snRNA-seq on healthy (*n*=9), APAP-ALF explant (*n*=10), and NAE-ALF explant (*n*=12) human liver tissue, including paired snRNA-seq and ST datasets from select patients (SI Table 1). This combined snRNA-seq dataset (72,262 nuclei) was annotated using signatures of known lineage markers (Fig. 1m,n, Extended Data Fig. 1e-i, Extended Data Fig. 2a-d, SI Table 2-4). Isolating and clustering the healthy hepatocytes from this dataset, and applying SPATA-derived zonal signatures, we found consensus between the ST and snRNA-seq approaches (Extended Data Fig. 2e-g).

We provide an open-access, interactive browser to allow assessment and visualisation of gene expression in multiple hepatic cell lineages in our healthy, APAP-ALF, and NAE-ALF ST and snRNA-seq datasets (https://shiny.igc.ed.ac.uk/7efa5350ba94425388e47ea7cdd5aa64/).

### Uncovering migratory hepatocytes in liver regeneration

Hepatocyte replenishment is a key process during liver regeneration. Given this, we focussed on hepatocytes from the annotated human snRNA-seq dataset (Fig. 2a). In line with multiplex smFISH, ST, and IF-based quantitation (Fig. 1a,i-k), we observed disruption of zonation and a robust proliferative transcriptomic response in hepatocytes following APAP-ALF and NAE-ALF compared to healthy controls (Fig. 2b, Extended Data Fig. 3a). Clustering these human hepatocytes uncovered a distinct ANXA2^+^ subpopulation in APAP-ALF and NAE-ALF compared to healthy controls (Fig. 2a,b, Extended Data Fig. 3b,c, SI Table 3).

**Figure 2:**
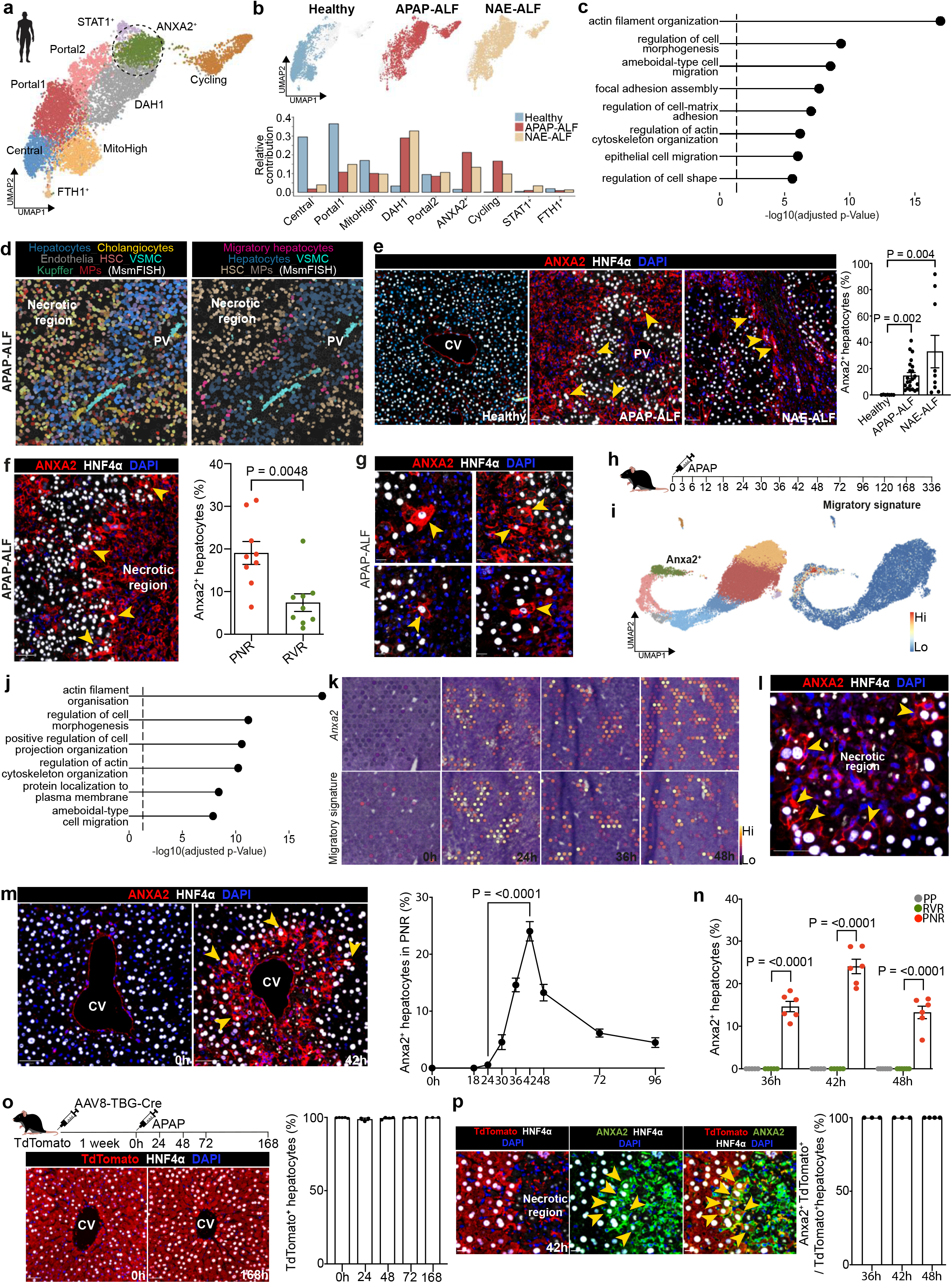
Uncovering migratory hepatocytes in liver regeneration. **a**, UMAP of hepatocyte nuclei from healthy, APAP-ALF, and NAE-ALF human liver explants, coloured by cluster. DAH1 = Disease Associated Hepatocytes 1. **b**, UMAPs (top) and barplots (bottom) displaying relative contribution of healthy, APAP-ALF, and NAE-ALF samples to each hepatocyte cluster. DAH1 = Disease Associated Hepatocytes 1. **c**, GO terms enriched in human ANXA2+ hepatocytes. **d**, Multiplex smFISH of cell types in APAP-ALF human liver (left). Multiplex smFISH showing migratory hepatocytes in APAP-ALF human liver in relation to other cell lineages (right). Gene modules in SI Table 2. **e**, Representative immunofluorescence images of ANXA2 (red), HNF4α (hepatocytes, white) and DAPI (nuclear stain, blue) in healthy, APAP-ALF, and NAE-ALF human liver tissue (left, scale bar 50µm). Percentage ANXA2+ hepatocytes in healthy, APAP-ALF, and NAE-ALF human livers (right), n=7 (healthy), n=22 (APAP-ALF), n=9 (NAE-ALF). Unpaired t-test. Data are mean±SEM. **f**, Representative immunofluorescence images of ANXA2 (red), HNF4α (hepatocytes, white) and DAPI (nuclear stain, blue) in APAP-ALF human liver (left). Yellow arrowheads denote ANXA2+ hepatocytes. Scale bar 50µm. Percentage ANXA2+ hepatocytes present in PNR and RVR of APAP-ALF human liver (right, n=9). Paired t-test, t=3.86, df=8. Data are mean±SEM. **g**, Representative immunofluorescence images of ANXA2 (red), HNF4α (hepatocytes, white) and DAPI (nuclear stain, blue) in APAP-ALF human liver (n=4). Yellow arrowheads denote ANXA2+ hepatocytes with migratory morphology. Scale bar 20µm. **h**, Schematic of time points processed for snRNA-seq post APAP-induced liver injury in mice. **i**, UMAP of mouse hepatocyte nuclei from all timepoints post APAP-induced acute liver injury, coloured by cluster (left). Application of human migratory hepatocyte gene module to mouse hepatocytes, showing corresponding region enriched in migratory gene signature (right). **j**, GO terms enriched in mouse migratory hepatocytes. **k**, Spatial transcriptomic expression of *Anxa2* (top) and migratory gene signature (bottom) in mouse liver post APAP-induced acute liver injury. **l**, Representative immunofluorescence images of ANXA2 (red), HNF4α (hepatocytes, white), DAPI (nuclear stain, blue) in mouse liver at 42hrs post APAP-induced acute liver injury (left). Yellow arrowheads denote ANXA2+ hepatocytes, scale bar 50µm. **m**, Representative immunofluorescence images of ANXA2 (red), HNF4α (hepatocytes, white) and DAPI (nuclear stain, blue) in mouse liver post APAP-induced acute liver injury (left). CV, central vein. Scale bar 50µm. Percentage ANXA2+ hepatocytes across timepoints post APAP-induced acute liver injury in the peri-necrotic region (PNR, right). Two-way ANOVA, n=3 (0h, 18h), n=6 (24-96h), *F*=44.60, df=8,34. Data are mean±SEM. **n**, Percentage ANXA2+ hepatocytes by region at peak expression timepoints. PP, peri-portal; RVR, remnant viable region; PNR, peri-necrotic region. Two-way ANOVA, n=6, *F*=15.85, df=2,15. Data are mean±SEM. **o**, Schematic depicting experimental protocol for lineage tracing of hepatocytes in AAV8.TBG.Cre-activated *R26LSLtdTomato* mice post APAP-induced liver injury (top). Representative immunofluorescence images of hepatocytes (tdTomato, red), HNF4α (hepatocytes, white) in select timepoints post APAP-induced acute liver injury (left, scale bar 50µm). TdTomato+ hepatocytes (HNF4α+) as a percentage of all hepatocytes post APAP-induced liver injury. One-way ANOVA, n=5 (0h), n=3 (24h, 72h, 168h), n=4 (48h), *F*=1.61, df=4,13. Data are mean±SEM. **p**, Representative immunofluorescence images of hepatocytes (tdTomato, red), ANXA2 (green), HNF4α (hepatocytes, white) in AAV8.TBG.Cre-activated *R26LSLtdTomato* mice 42hrs post APAP-induced liver injury (left). Yellow arrowheads denote ANXA2+TdTomato+ hepatocytes. Scale bar 20µm. Anxa2+TdTomato+ hepatocytes (HNF4α+) as a percentage of all TdTomato+ hepatocytes at peak (ANXA2+ hepatocyte) timepoints post APAP-induced liver injury (right). n=3 (36h, 42h), n=4 (48h). Data are mean±SEM.

GO analysis of the ANXA2^+^ hepatocytes (Fig. 2c, SI Table 4) highlighted terms including ameboidal-type cell migration, regulation of cell morphogenesis, epithelial cell migration, and regulation of cell shape, suggesting a migratory cell phenotype. Differentially expressed genes in the ANXA2^+^ hepatocyte subpopulation allowed us to define a migratory hepatocyte signature (Extended Data Fig. 3d, SI Table 2) which, when applied to ST, was observed in and around the necrotic region in APAP-ALF and NAE-ALF (Extended Data Fig. 3e). Multiplex smFISH enabled delineation of multiple cell lineages (Fig. 2d, left; Extended Data Fig. 3f) and demonstrated expression of a migratory hepatocyte signature in a subpopulation of hepatocytes adjacent to the necrotic region in APAP-ALF, which was absent in healthy liver tissue (Fig. 2d, right; Extended Data Fig. 3g, SI Table 2).

IF staining confirmed significantly increased ANXA2^+^ hepatocytes in APAP-ALF and NAE-ALF livers compared to healthy controls (Fig. 2e), and increased ANXA2^+^ hepatocytes in the PNR in APAP-ALF (Fig. 2f). ANXA2^+^ hepatocytes exhibited a motile morphology with ruffled membranes and extending lamellipodia, characteristic of migratory cells (Fig. 2g). This ANXA2^+^ hepatocyte subpopulation was also observed in other causes of human ALF including hepatitis A and B, and other drug-induced liver injuries (Extended Data Fig. 3h).

To determine if a corollary migratory hepatocyte subpopulation exists in a mouse model of APAP-induced acute liver injury, we performed snRNA-seq (59,051 nuclei) and ST of mouse liver across multiple timepoints (Fig. 2h, Extended Data Fig. 4a-d). Applying the human migratory hepatocyte signature to mouse hepatocytes (Fig. 2i, SI Video 1) identified an analogous ANXA2^+^ subpopulation (Extended Data Fig. 4e) emerging in response to APAP-induced mouse liver injury, exhibiting comparable migratory ontology (Fig. 2j, SI Video 2). Akin to our findings in human APAP-ALF, spatio-temporal profiling revealed disruption of hepatocyte zonation (Extended Data Fig. 4f) in APAP-induced mouse liver injury. Applying SPATA-derived mouse zonal signatures to the hepatocyte subpopulations demonstrated consensus between the ST and snRNA-seq datasets (SI Video 3, SI Table 2). Further, ST delineated the migratory hepatocyte subpopulation around the necrotic region following APAP-induced mouse liver injury (Fig. 2k). IF staining confirmed the presence of ANXA2^+^ hepatocytes following APAP-induced mouse liver injury, and these hepatocytes exhibited a similar morphology to the ANXA^+^ hepatocytes observed in human APAP-ALF, with ruffled membranes and lamellopodia (Fig. 2l). ANXA2^+^ hepatocytes increased following APAP-induced mouse liver injury, peaking at 42 hours (Fig. 2m), and were specifically enriched in the PNR (Fig. 2n).

To determine whether new hepatocytes derive from hepatocytes following APAP-induced mouse liver injury, we lineage-traced hepatocytes using adeno-associated viral AAV8.TBG.Cre injected into *R26R*^*LSL*^*tdTomato* mice. AAV8.TBG.Cre injection activated tdTomato (tdTom) expression in 99.8% (±0.1%SEM) of hepatocytes (HNF4α^+^) in healthy mouse liver (Fig. 2o). Hepatocyte lineage tracing following APAP-induced mouse liver injury demonstrated that 99.9% (±0.1%SEM; day 7) of new hepatocytes derived from tdTom^+^ hepatocytes (Fig. 2o). Furthermore, 100% of ANXA2^+^ hepatocytes derived from tdTom^+^ hepatocytes at all timepoints studied (Fig. 2p).

### Wound closure precedes hepatocyte proliferation during APAP-induced liver regeneration

Investigating the dynamics of liver regeneration following APAP-induced mouse liver injury, we uncovered a temporal disconnect between wound closure (as assessed by percentage necrotic area) and hepatocyte proliferation. Peak hepatocyte necrosis occurred at 30h post APAP-induced liver injury (22.3%±1.3%SEM), with percentage necrotic area decreasing by 30.9% at 42h (15.4%±1.4%SEM) and by 58.3% at 48h (9.3%±1%SEM) (Fig. 3a). Wound closure preceded the onset of hepatocyte proliferation, which peaked at 72h post APAP-induced liver injury (Fig. 3a).

**Figure 3:**
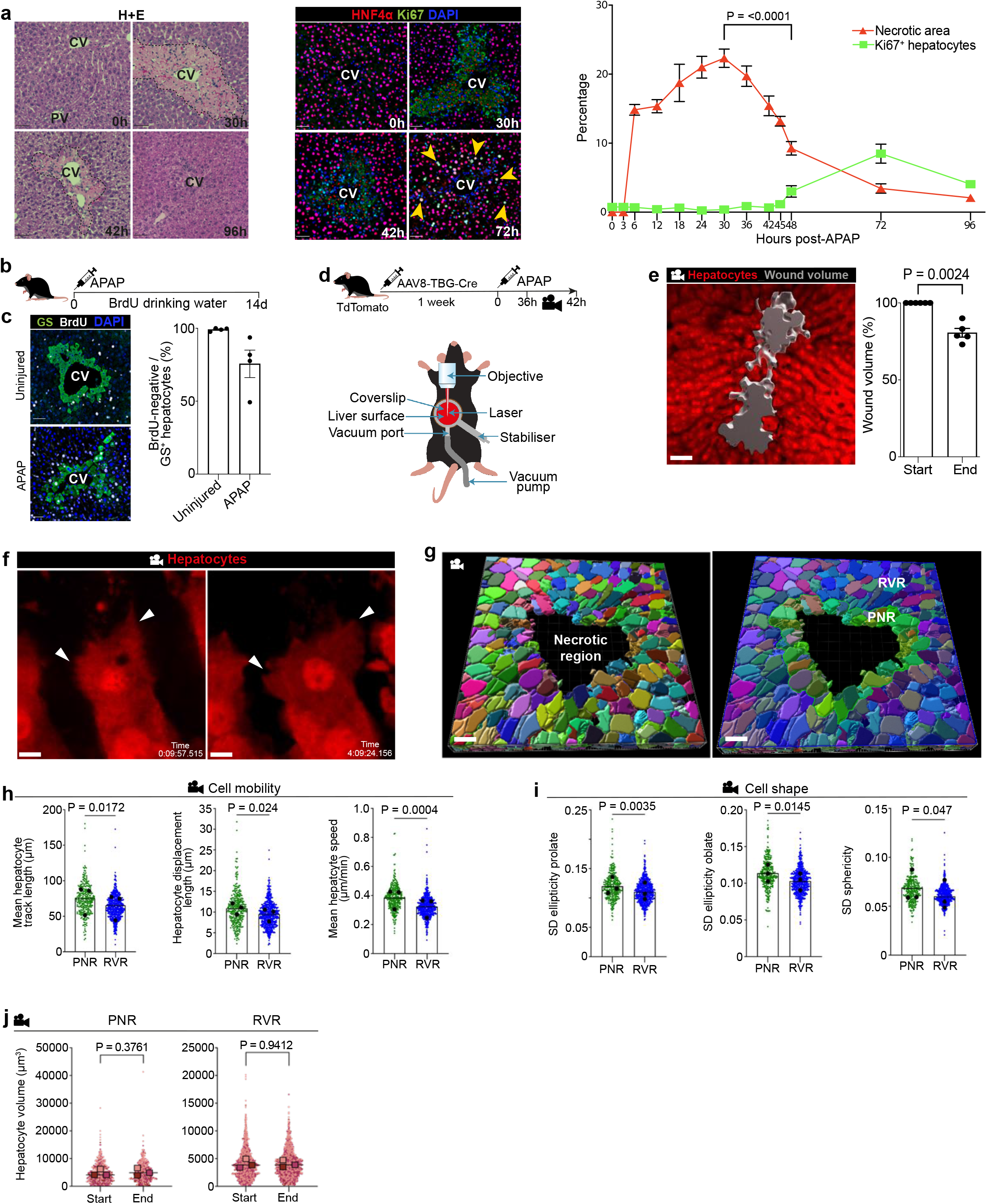
Hepatocyte migration and wound closure precedes hepatocyte proliferation during APAP-induced liver regeneration. **a**, Representative haematoxylin and eosin staining across select time points following APAP-induced mouse liver injury (left). Representative immunofluorescence staining: Ki67 (green), HNF4α (hepatocytes, red), DAPI (nuclear stain, blue) across select time points following APAP-induced mouse liver injury (middle). Scale bar 50µm. Quantification of necrotic area (red) and hepatocyte proliferation (green) following APAP-induced mouse liver injury (right). Two-way ANOVA, n=3 (0-18h), n=8 (24-96h), *F*=39.80, df=12,57. Data are mean±SEM. **b**, Schematic depicting BrdU dosing of mice post APAP-induced acute liver injury (top). **c**, Representative immunofluoresence staining (left) and quantification (right) of percentage BrdU-negative GS+ hepatocytes adjacent to the central vein: GS (green), BrdU (white), DAPI (nuclear stain, blue) Scale bar 50µm. n=4. Data are mean±SEM. **d**, Schematic depicting experimental protocol (top) and intravital microscopy (IVM) setup (bottom). **e**, Representative IVM snapshot of APAP-induced mouse liver injury with volume rendering of wound area (left). Scale bar 50µm. Quantification of change in wound volume at start and end of IVM imaging session (right). Paired t-test, n=5, t=6.82, df=4. Data are mean±SEM. **f**, Representative IVM snapshots of motile hepatocytes, white arrowheads marking areas of membrane ruffling / lamellipodia formation. Scale bar 5µm. **g**, Representative IVM images of Cellpose-segmented hepatocytes (left) and binning of hepatocytes into peri-necrotic (PNR, green) or remnant viable (RVR, blue) regions (right). Scale bar 40µm. **h**, Quantification of cell mobility, mean track length (left, paired t-test, t=7.52, df=2), mean displacement length (middle, paired t-test, t=6.34, df=2) and mean speed (right, paired t-test, t=47.20, df=2) in PNR (green) or RVR (blue) hepatocytes. n=3 mice. Data are mean. **i**, Quantification of cell shape changes over time. Ellipticity prolate (left, paired t-test, t=16.79, df=2). Ellipticity oblate (middle, paired t-test, t=8.21, df=2). Sphericity (right, paired t-test, t=4.45, df=2). SD, standard deviation. n=3 mice. Data are mean. **j**, Quantification of hepatocyte volume at start and end of IVM, for each mouse. Paired t-test, n=2 (mouse 1), n=3 (mouse 2,3) regions per mouse. Data are mean±SEM.

To assess whether hepatocyte repopulation of the necrotic area immediately adjacent to the central vein is driven by proliferation, we gave BrdU in drinking water to label all proliferating hepatocytes following APAP-induced mouse liver injury (Fig. 3b). Hepatocyte proliferation had returned to baseline levels by day 14, with complete wound closure (Extended Data Fig. 5a). Glutamate synthetase (GS) is expressed exclusively in hepatocytes adjacent to the central vein (Fig. 3c), and 75.6% (±9.4%SEM) of GS-positive hepatocytes were BrdU-negative 14 days post APAP-induced liver injury, implying that the majority of hepatocytes originate from pre-existing hepatocytes outwith the necrotic area, which migrate adjacent to the CV (Fig. 3c).

Having identified a migratory hepatocyte subpopulation in human and mouse APAP-induced liver injury (Fig. 2), and given that hepatocyte proliferation is not the major contributor to wound closure (Fig. 3a,c), we performed 4-D intravital microscopy (IVM) to investigate whether hepatocyte migration occurs *in vivo* following APAP-induced liver injury (Fig. 3d). We used AAV8.TBG.Cre combined with two fluorescent mouse reporter lines to label hepatocytes: *Hep;tdTom* (single-fluorescent reporter mice that express cytoplasmic tdTomato after Cre-mediated recombination) and *Hep;mGFP* (double-fluorescent reporter mice expressing membrane-targeted GFP after Cre-mediated excision). IVM did not affect levels of hepatocyte necrosis, proliferation, or hepatocyte ANXA2 expression compared to non-IVM imaged mice at 42hrs post APAP-induced liver injury (Extended Data Fig. 5b-d). 4-D IVM of *Hep;tdTom* reporter mice demonstrated centrilobular hepatocyte necrosis in real-time during APAP-induced mouse liver injury (SI Video 4,5).

Due to the prevalence of ANXA2^+^ hepatocytes (Fig. 2k,l) and wound closure activity (Fig. 3a) between 36 to 42 hours post APAP-induced mouse liver injury, we performed IVM during this timeframe (Fig. 3e). 4-D IVM of *Hep;tdTom* mice demonstrated collective migration of the hepatocyte sheet (SI Video 6a,b). Rendering of wound volume (Fig. 3e, left) showed 19.4% (±2.8%SEM) reduction in necrosis during the imaging period (Fig. 3e, right). Intravital imaging of *Hep;tdTom* mice identified hepatocytes with a motile morphology, including membrane ruffling and the formation of lamellipodia at the hepatocyte leading edge abutting the wound (Fig. 3f, SI Videos 6-13). Using *Hep;mGFP* to clearly segment individual hepatocytes (Fig. 3g, left), we then classified hepatocytes into the PNR (green) or the RVR (blue) to compare cell mobility and shape between the two regions (Fig. 3g, right). Measures of hepatocyte mobility (mean track length, displacement length, and speed) were greater in hepatocytes in the PNR compared to hepatocytes in the RVR (Fig. 3h). Assessment of cell shape over time demonstrated that hepatocytes in the PNR displayed greater deviation in ellipticity (prolate and oblate) and sphericity compared to hepatocytes in the RVR (Fig. 3i). During the imaging period, there was no change in hepatocyte volume in either the PNR or RVR (Fig. 3j). Taken together, these data demonstrate that hepatocyte migration is a major mechanism of wound closure following APAP-induced liver injury, and that wound closure is not mediated by hepatocyte hypertrophy.

### Hepatocyte ANXA2 regulates hepatocyte migration and wound closure during APAP-induced liver regeneration

ANXA2, which we have identified as a key marker in both human and mouse migratory hepatocytes, has previously been shown to regulate cell migration in other disease settings including carcinogenesis^16-18^. Knockdown of ANXA2 in a human hepatocellular carcinoma cell line (Huh7) decreased wound closure at 72h in a scratch wound assay (Fig. 4a, Extended Data Fig. 6a). Primary hepatocytes from uninjured mouse livers increased *Anxa2* gene expression in response to plating on tissue culture plastic (Fig. 4b). Using this *in vitro* model system, inhibition of *Anxa2* expression in HGF (hepatocyte growth factor)-stimulated primary mouse hepatocytes (Extended Data Fig. 6b) reduced wound closure (Fig. 4c), and this effect was not mediated by a reduction in hepatocyte proliferation (Fig. 4d). *Met* (HGF receptor) gene expression was similar between Scrmb-siRNA and *Anxa2*-siRNA treated hepatocytes (Fig. 4e).

**Figure 4:**
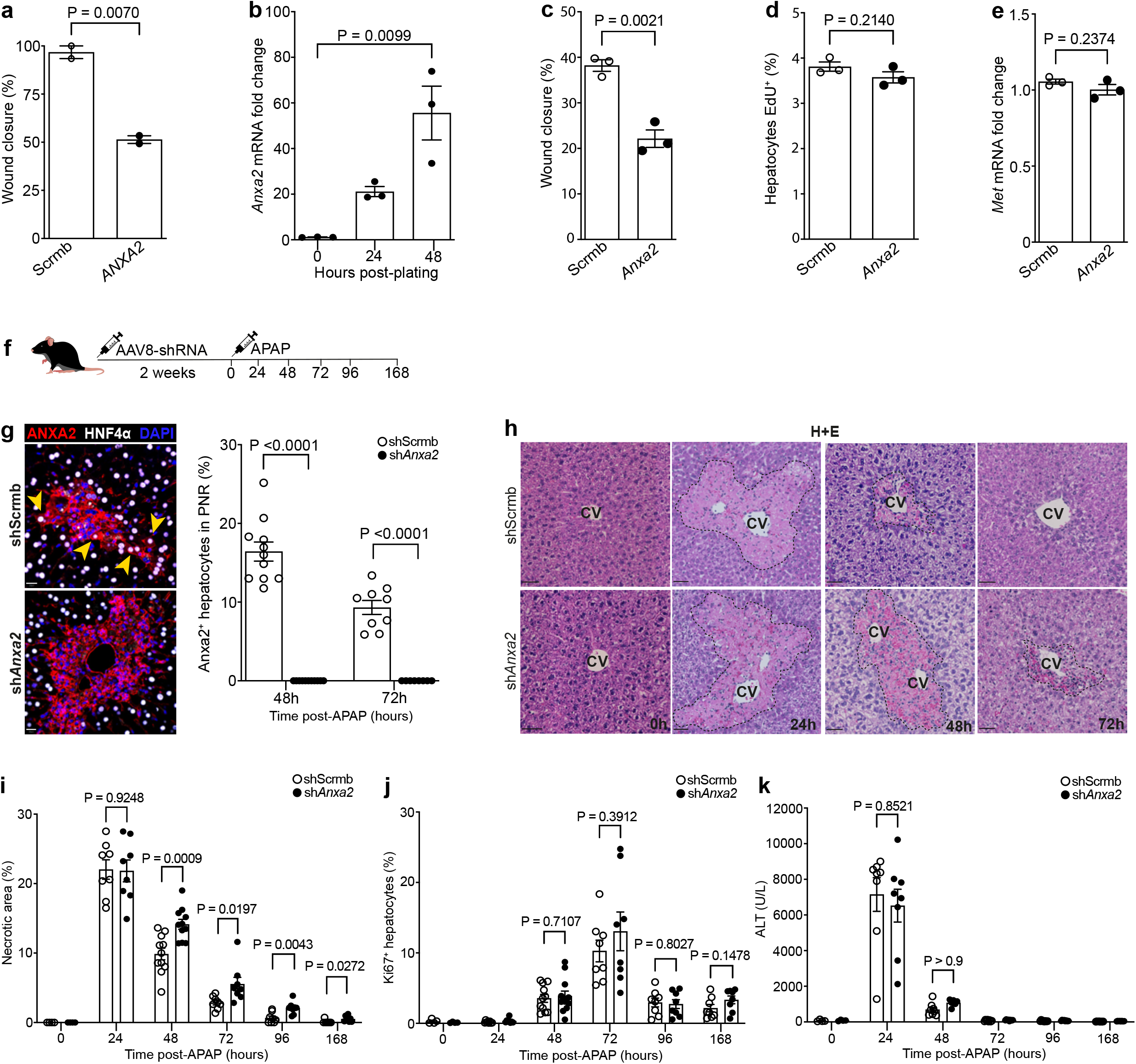
Hepatocyte ANXA2 regulates hepatocyte migration and wound closure during APAP-induced liver regeneration. **a**, Percent coverage of scratch wound area 72hrs post-wounding in Huh7 human hepatocyte cell line, in control (Scrmb) compared to ANXA2-siRNA treated cells. Unpaired t-test, t=11.89, df=2, two independent experiments. Data are mean±SEM. **b**, Timecourse of *Anxa2* gene expression (RT-qPCR analysis) in plated primary mouse hepatocytes. Unpaired t-test, t=4.62, df=4, three independent experiments. Data are mean±SEM. **c**, Percent coverage of scratch wound area 72hrs post-wounding of primary mouse hepatocytes, in control (Scrmb) compared to *Anxa2*-siRNA treated cells. Unpaired t-test, t=7.04, df=4, three independent experiments. Data are mean±SEM. **d**, Percent EdU+ hepatocytes 72hrs post-wounding, in control (Scrmb) compared to *Anxa2*-siRNA treated cells. Unpaired t-test, t=1.48, df=4, three independent experiments. Data are mean±SEM. **e**, *Met* gene expression (RT-qPCR analysis) in primary mouse hepatocytes, control (Scrmb) compared to *Anxa2*-siRNA treated cells. Unpaired t-test, t=1.39, df=4, three independent experiments. Data are mean±SEM. **f**, Schematic depicting experimental protocol for *in vivo* hepatocyte *Anxa2* knockdown and APAP-induced mouse liver injury. **g**, Representative immunofluorescence staining of mouse liver from AAV8-shScrmb or AAV8-sh*Anxa2* treated mice at 48 hours post APAP-induced acute liver injury (left). ANXA2 (red), HNF4α (hepatocytes, white) DAPI (nuclear stain, blue). Scale bar 20µm. Percentage ANXA2+ hepatocytes post APAP-induced acute liver injury in the peri-necrotic region (PNR) of AAV8-shScrmb or AAV8-sh*Anxa2* treated mice (right). Two-way ANOVA, n=11 (48h), n=9 (72h), *F*=20.64, df=1. Data are mean±SEM. **h**, Representative haematoxylin and eosin staining of mouse livers from AAV8-shScrmb or AAV8-sh*Anxa2* treated mice across timepoints post APAP-induced acute liver injury. Scale bar 50µm. **i**, Quantification of necrotic area following APAP-induced acute liver injury in AAV8-shScrmb or AAV8-sh*Anxa2* treated mice. Unpaired t-test, n=4 (0h), n=8 (24h), n=11 (48h), n=8 (72-168h). Data are mean±SEM. **j**, Quantification of Ki67+ hepatocytes following APAP-induced acute liver injury in AAV8-shScrmb or AAV8-sh*Anxa2* treated mice. Unpaired t-test, n=4 (0h), n=8 (24h), n=11 (48h), n=8 (72-168h). Data are mean±SEM. **k**, Alanine transaminase (ALT) following APAP-induced acute liver injury in AAV8-shScrmb or AAV8-sh*Anxa2* treated mice. Unpaired t-test, n=4 (0h), n=8 (24-168h). Data are mean±SEM.

To determine the functional role of hepatocyte ANXA2 during APAP-induced mouse liver injury, we used AAV8-shRNA-*Anxa2* to knockdown *Anxa2* specifically in hepatocytes *in vivo* (Fig. 4f,g). Knockdown of hepatocyte *Anxa2* abrogated wound closure compared to control (AAV8-shRNA-Scrmb) following APAP-induced mouse liver injury (Fig. 4h,i); this effect was not mediated by a reduction in hepatocyte proliferation, or by ongoing hepatocyte injury (Fig. 4j,k). *Anxa2* gene expression was also observed in leukocytes, mesenchymal and endothelial cells during APAP-induced mouse liver injury (Extended Data Fig. 6c). Treatment with AAV8-shRNA-*Anxa2* did not affect *Anxa2* expression in leukocytes (CD45^+^), mesenchymal and endothelial cells during APAP-induced mouse liver injury compared to control (Extended Data Fig. 6d-f). Furthermore, total numbers of leukocytes, mesenchymal and endothelial cells were similar between AAV8-shRNA-Scrmb and AAV8-shRNA-*Anxa2* treated groups (Extended Data Fig. 6g). In summary, these data demonstrate that hepatocyte ANXA2 expression regulates hepatocyte migration and wound closure during APAP-induced liver regeneration.

## Discussion

To advance our understanding of human liver regeneration and to help inform design of pro-regenerative therapies, we generated the first single-cell, pan-lineage atlas of human liver regeneration. We provide an open-access, interactive browser to allow assessment and visualisation of gene expression in all hepatic cell lineages (snRNA-seq and ST datasets) in healthy and regenerating human and mouse liver (https://shiny.igc.ed.ac.uk/7efa5350ba94425388e47ea7cdd5aa64/). Using a cross-species, integrative multimodal approach, we uncovered a novel migratory hepatocyte subpopulation which is critical in mediating successful wound healing and reconstitution of normal hepatic architecture following liver injury.

Hepatocyte proliferation has previously been considered a major driver of hepatic wound closure following necro-inflammatory liver injury^2,19^. We observed a substantial hepatocyte proliferative response in human APAP-ALF explant livers compared to uninjured human liver, demonstrating that hepatocyte proliferation is robust and relatively unimpeded in human APAP-ALF. However, despite vigorous hepatocyte proliferation, the necrotic wound area in human APAP-ALF explant livers remained substantial at time of transplantation. These data suggest that mechanisms and processes other than hepatocyte proliferation are critical to effect successful wound closure following human liver injury.

Furthermore, previous studies in transgenic mice have suggested that hepatocyte proliferation is not a key mechanism regulating wound closure following necro-inflammatory liver injury. Plasminogen-knockout mice displayed persistent centrilobular damage and severe impairment of repair following carbon tetrachloride (CCl^4^)-induced acute liver injury, despite a normal hepatocyte proliferative response compared to control mice^20^. Conditional knockout of the HGF receptor (c-Met) in mouse hepatocytes followed by CCl^4^-induced acute liver injury impaired centrilobular wound closure and restitution of normal tissue architecture, despite similar levels of hepatocyte proliferation compared to controls^21^.

Following our discovery of a novel migratory hepatocyte subpopulation during human and mouse liver regeneration, we used the mouse model of APAP-induced liver injury to investigate the dynamics of liver regeneration. We uncovered a temporal disconnect between centrilobular wound closure and hepatocyte proliferation, demonstrating that wound closure precedes hepatocyte proliferation following APAP-induced mouse liver injury. Furthermore, continual administration of BrdU to label all hepatocytes following APAP-induced mouse liver injury demonstrated that the majority of hepatocytes in the area immediately adjacent to the central vein did not arise from hepatocyte proliferation. 4-D intravital imaging of APAP-induced mouse liver injury identified motile hepatocytes, displaying membrane ruffling and the formation of dynamic protrusions at the leading edge, with collective cell migration of the hepatocyte sheet to effect wound closure. Depletion of hepatocyte ANXA2 expression reduced HGF-induced migration of human and mouse hepatocytes *in vitro*, and depletion of hepatocyte ANXA2 *in vivo* abrogated necrotic wound closure following APAP-induced liver injury in mice.

Wound healing in the skin is largely driven by keratinocyte migration^22^, where rapid reconstitution of the epidermal barrier is vital to stop invasion by pathogens. Our findings point towards a similar mechanism occurring in the liver. Expeditious wound closure in the liver, with restoration of the gut-liver barrier following acute, massive epithelial injury, may be a key determinant of patient outcome in ALF where sepsis and multi-organ failure are the commonest cause of death^3-5^. Recent studies in mice have demonstrated that APAP-induced liver injury causes impairment of intestinal barrier integrity and increased bacterial translocation^23-25^, highlighting that APAP toxicity causes barrier breakdown in both the gut and the liver. We propose that the liver prioritises rapid re-epithelialisation via migration of the hepatocyte sheet before hepatocyte proliferation, as rapid restoration of the gut-liver barrier is paramount to survival by preventing bacterial dissemination with subsequent sepsis and multi-organ failure.

In summary, our work dissects unanticipated aspects of human liver regeneration, uncovering a novel migratory hepatocyte subpopulation which mediates wound closure following liver injury. Therapies designed to promote hepatocyte migration with rapid reconstitution of normal hepatic microarchitecture and reparation of the gut-liver barrier may open up a new area of therapeutic discovery in regenerative medicine.

## Methods

### Study subjects

#### University of Edinburgh, UK

Local approval for procuring human liver tissue for single-nuclei RNA sequencing, spatial transcriptomics and histological analysis was obtained from the Scotland ‘A’ Research and Ethics Committee (16/SS/0136) and the NRS BioResource and Tissue Governance Unit (study number SR574), following review at the East of Scotland Research Ethics Service (reference 15/ES/0094). Written informed consent was obtained from the subject or a legally authorised representative prior to enrolment per local regulations. Acute liver failure liver tissue was obtained intraoperatively from patients undergoing orthotopic liver transplantation at the Scottish Liver Transplant Unit, Royal Infirmary of Edinburgh. Patient demographics are summarized in SI Table 1 for patients transplanted for APAP-induced ALF and nonA-E ALF. Healthy background non-lesional liver tissue was obtained intraoperatively from patients undergoing surgical liver resection for solitary colorectal metastasis at the Hepatobiliary and Pancreatic Unit, Department of Clinical Surgery, Royal Infirmary of Edinburgh. Patients with a known history of chronic liver disease, abnormal liver function tests or those who had received systemic chemotherapy within the last four months were excluded from this cohort. For histological assessment of human ALF and chronic liver disease tissue, anonymized unstained formalin-fixed paraffin-embedded liver biopsy sections were provided by the Lothian NRS Human Annotated Bioresource under authority from the East of Scotland Research Ethics Service REC 1, reference 15/ES/0094.

#### United States Acute Liver Failure Study Group (ALFSG) network

This consortium of U.S. liver centers was established in 1998 to better define causes and outcomes of acute liver injury and ALF. The study protocol was approved by the local institutional review boards of the participating sites. Written informed consent was obtained from the subject or a legally authorized representative prior to enrolment per local regulations. Sites obtained portions of fresh explanted liver tissue cut into 1cm^3^ pieces, placed into individual cryovials and stored at −80 °C until requested for study. The ALFSG was supported by the National Institute of Diabetes and Digestive and Kidney Diseases (NIDDK; grant no.: U‐01‐58369). The samples used in this study were supplied by the NIDDK Central Repositories. This article does not necessarily reflect the opinions or views of the NIDDK Central Repositories or the NIDDK.

#### University of Cambridge, UK

Patients were recruited at Addenbrooke’s Hospital, Cambridge, UK with approval of the Local Research Ethics Committee (16/NI/0196 & 20/NI/0109). Written informed consent was obtained from the subject or a legally authorised representative prior to enrolment per local regulations. Liver tissue from patients with ALF was derived from explanted livers at the time of transplantation. All tissue samples were snap-frozen in liquid nitrogen and stored at −80 °C in the Human Research Tissue Bank of the Cambridge University Hospitals NHS Foundation Trust.

#### University of Birmingham, UK

Human liver tissue obtained from the University of Birmingham, UK was obtained under local ethical approval (reference 06/Q2708/11, 06/Q2702/61) and written informed consent was obtained from the subject or a legally authorised representative prior to enrolment per local regulations. Liver tissue was acquired from explanted livers from patients undergoing orthotopic liver transplantation at the Queen Elizabeth Hospital, Birmingham. All tissue samples were snap-frozen in liquid nitrogen and stored at −80 °C before being processed and shipped by the Birmingham Human Biomaterials Resource Centre (reference 09/H1010/75; 18-319).

### Mice

Mice used for all experiments were male, 8–12 weeks old and housed under pathogen–free conditions at the University of Edinburgh. All experiments were performed in accordance with the UK Home Office Regulations. C57BL/6JCrl mice were obtained from Charles River Laboratories (UK). mTmG (Jax 007676; *B6*.*129(Cg)-Gt(ROSA)26Sor*^*tm4(ACTB-tdTomato,-EGFP)Luo*^*/J*)^26^ and TdTomato (Jax 007914; *B6*.*Cg-Gt(ROSA)26Sor*^*tm14(CAG-tdTomato)Hze*^*/J*)^27^ reporter mice were obtained from Jackson Laboratories. For the acetaminophen-induced acute liver injury model, mice were fasted for 12 hours prior to intraperitoneal injection (IP) with 300 mg/kg of acetaminophen dissolved in sterile phosphate-buffered saline (PBS) as described previously^28^. For hepatocyte-specific AAV8-Cre mediated reporter gene induction, stock AAV8.TBG.PI.Cre.rBG (AAV8-TBG-Cre; a gift from James M. Wilson (Addgene plasmid #107787); stored at -80°C) was thawed on ice, diluted in sterile PBS to achieve a working titre of 2×10^12 genetic copies (GC)/ml and was subsequently stored at -20°C until usage. On the day of injection the diluted AAV was thawed and each mouse was injected via the tail vein with 100μl (2×10^11 GC/mouse)^29^. Mice were left for one week prior to acetaminophen-induced acute liver injury. For *in vivo* hepatocyte-specific knockdown of *Anxa2*, mice were intravenously injected with 1×10^12 GC/ml AAV8-GFP-U6-mANXA2-shRNA (sh*Anxa2*) or AAV8-GFP-U6-scrmb-shRNA (shScrmb) and left for two weeks prior to acetaminophen-induced acute liver injury. For administration of 5-Bromo-2’-deoxyuridine (BrdU) in drinking water, BrdU was dissolved in drinking water at a concentration of 0.8mg/ml.

### Nuclei isolation for single-nuclei RNA sequencing (snRNA-seq)

Human and mouse liver for snRNA-seq was processed as described previously using the TST method^30^. Mouse liver nuclei isolation was performed on n=3 mice per timepoint and nuclei were pooled for snRNA-seq.

### Droplet-based snRNA-seq

Single nuclei were processed through the 10X Genomics Chromium Platform using the Chromium Single Cell 3′ Library and Gel Bead Kit v3 (10X Genomics, PN-1000075) and the Chromium Single Cell B Chip Kit (10X Genomics, PN-1000074) as per the manufacturer’s protocol, and as previously described^31^. Libraries were sequenced on either an Illumina HiSeq 4000 or NovaSeq 6000.

### Spatial transcriptomics

Unfixed liver tissues were embedded in Tissue-Tek (OCT) and snap-frozen. Samples were then cryosectioned (10µm) and placed on pre-chilled Visium (10X Genomics) tissue optimization slides or Visium spatial gene expression slides. Spot size is 55µm, with 100µm between spot centroids. Tissue sections were processed as per the manufacturers protocol. Based on optimisation time course experiments, tissue was permeabilised for 18 minutes.

### Multiplex smFISH

Unfixed snap-frozen liver tissues were cryosectioned (10µm) onto Resolve Biosciences slides and sent on dry ice to Resolve Biosciences for processing. Gene probes were designed using Resolve Biosciences proprietary design algorithm. Probe details are included in the key resources table. Following sample imaging, spot segmentation, pre-processing and signal segmentation and decoding, final image analysis was performed in R programming language.

### Liver Intravital Microscopy (Liver-IVM)

Single colour tdTomato imaging was performed using a Discovery (Coherent) ‘multiphoton’ laser tuned to 1050 nm through a 20x VIS-IR corrected water immersion objective (N.A. 1.0) by placing a water drop on top of the coverslip. Dual colour eGFP/tdTomato imaging was performed using the same set up with the laser tuned to 1000 nm. Non-descanned GaAsP detectors (GFP NDD filter BP 500-550nm; tdTomato NDD Filter BP 575-610nm) were used to initially obtain an overview image after which three peri-central vein fields were selected for time-lapse imaging. These three fields were then imaged as z-stacks (40-50µm) every 10 minutes for 6 hours. Following administration of acetaminophen (350mg/kg) mice were anaesthetised with isoflurane (4% induction, 1-1.5% maintenance) in approximately 95% oxygen (0.8 L / min.) produced by an oxygen scavenger (Vettech). Coat above the liver was clipped back, Lacrilube applied to the eyes, mice were then placed in a dorsal position on a heated stage on an upright LSM 880 NLO multiphoton microscope (Zeiss). An abdominal incision was made, exposing the surface of the liver, this was then stabilised using a custom coverslip-holding imaging vacuum stabilisation armature attached to the stage. The gentlest possible vacuum was applied to the surface in contact with the liver, holding it in place against the coverslip. Mice received sub-cutaneous fluids every 45 minutes during imaging. At the end of the imaging session mice were humanely killed under general anaesthesia by cervical dislocation.

### Liver-IVM processing and segmentation

Timelapse image analysis and visualization was performed using Imaris 9.7 (Bitplane). Imaris ‘reference frame’ was firstly used to correct for xyz drift. To create 4D rendering of the wound the hepatocyte channel was inverted and smoothed using a gaussian filter. Imaris ‘surface’ tool was then used to create a surface corresponding to the wound. Object statistics were then exported to analyse the evolution of the surface’s volume over time (expressed as % of initial volume). To create 4D rendering of individual hepatocytes from the mTmG mice, registered images were first imported in Google drive and the online platform ZerocostDL4mic^32^ was used to perform Cellpose segmentation^33^ on the eGFP (hepatocytes) channel. Cellpose annotated images were processed in ImageJ using ‘Label to ROI’ plugin^34^ to create eroded ROIs followed by ‘Mask from ROI’ plugin to produce cell masks. In order to reduce non-specific segmentation, tdTomato signal was thresholded and substracted from the mask channel using the ‘channel arithmetics’ tool in Imaris. Imaris ‘surface’ tool was then used to create surfaces corresponding to the hepatocytes. Cell statistics were then exported to analyse morphodynamic parameters according to their relation to the wound (distance to the wound). Cell behaviour was determined using the track length and speed (indicating cell mobility) and the standard deviation of cell sphericity and ellipticity (indicating changes in cell shape over time).

### Immunohistochemistry and immunocytochemistry staining

Immunohistochemistry and immunofluorescence staining was performed on formalin-fixed, paraffin-embedded liver tissue sections (4µm). Slides were deparaffinized and immunofluorescently labelled using a Leica Bond RX_m_ automated robotic staining system. Antigen retrieval was performed using Bond Epitope Retrieval Solution 1 or 2 for 20 min in conjunction with heat application. Sections were then incubated with primary antibodies diluted in 0.1% Triton-X containing phosphate buffered saline (PBS). Sections were stained with DAPI (Sigma) and mounted on glass coverslips in ProLong Gold Antifade mounting medium and stored at 4°C until time of imaging. For *in vitro* EdU detection cells were washed in PBS then fixed for 10mins at room temperature with 4% formaldehyde solution in PBS, cells were then stained according to the manufacturer’s protocol. A full list of antibodies and conditions is included in SI Table 5.

### Histology image processing and segmentation

Slides were scanned using a Zeiss Axioscan Z1. For *in vitro* immunocytochemistry, wells were imaged using an EVOS FL Auto Imaging System. All image analysis was undertaken in QuPath v0.3.0^35^.

### Cell culture

Human immortalised hepatocyte cell line (Huh-7; Cell Lines Service 300156) was cultured using RPMI 1640 supplemented with 10% FBS and 2mM L-glutamine. Primary mouse hepatocytes were isolated and cultured as previously described^36^.

### Gene knockdown in hepatocytes

Gene knockdown in Huh7 and primary mouse hepatocytes was performed using siRNA. Cells were plated at 500,000 cells/ml (Huh7, Corning Costar; primary mouse and human hepatocytes Corning Primaria) followed by serum starvation overnight (in medium without FBS). siRNA duplexes with Lipofectamine RNAiMAX Transfection Reagent were prepared in OptiMEM according to the manufacturer’s recommendations and used at a concentration of 50nM. Cells were exposed to the duplex for 48 hours, in antibiotic-free media containing 2% FBS. Cells were harvested for RNA and RT-qPCR. Gene knockdown efficiency was assessed by RT-qPCR. Cells were treated with control siRNA (Qiagen, 1027280), siRNA for *ANXA2* (Human, Qiagen, Hs_ANXA2_8, SI02632385) or siRNA for *Anxa2* (Mouse, Qiagen, Mm_Anxa2_3, SI00167496).

### Scratch wound assay

Scratch wound assay was performed using the IncuCyte® system (Essen Bioscience). Huh7 cells were plated in IncuCyte® ImageLock Plates (Essen Bioscience, Ann Arbor, MI, USA) and treated as above for *ANXA2* gene knockdown. The sub-confluent monolayer was then wounded using the IncuCyte® Woundmaker. In order to obtain a confluent monolayer of primary mouse hepatocytes for wound assays, cells were plated as previously described^37^, with modifications. Briefly, three separate additions of 500,000 cells/ml were seeded onto Collagen I-coated IncuCyte® ImageLock plates at 2hr intervals. Non-adherent hepatocytes were removed between additions with warmed PBS. Cells were then treated as above for gene knockdown prior to wounding using the IncuCyte® Woundmaker. Following wounding cells were maintained in complete media with the addition of human Hepatocyte Growth Factor (HGF; 100ng/ml) and 10% FBS for the duration of the assay. EdU (10µM) was added to the media 24hrs prior to the end of the assay to assess proliferation. For analysis of wound healing the scratch wound plugin for the IncuCyte® Zoom was used. All experiments were performed as quadruple technical replicates, number of independent experiments are specified in figure legends.

### RNA extraction and RT–qPCR

RNA was extracted from primary mouse hepatocytes and Huh-7 cells using the RNeasy Plus Micro Kit and cDNA synthesis performed using the QuantiTect Reverse Transcription Kit according to the manufacturer’s protocol (Qiagen). Reactions were performed in triplicate in 384-well plate format. RT–qPCR was performed using PowerUp SYBR Green Master Mix, primers are detailed in SI Table 5. Samples were amplified on an ABI Quantstudio 5 (Applied Biosystems, Thermo Fisher Scientific). The 2−ΔΔCt quantification method, using GAPDH/Gapdh for normalization, was used to estimate the amount of target mRNA in samples, and expression calculated relative to average mRNA expression levels from control samples.

### Computational analysis

Four computational snRNA-seq datasets were analysed: 1) 72,262 human nuclei from healthy (n=9), APAP-ALF (n=10), and NAE-ALF (n=12) livers; 2) 59,051 mouse nuclei from an APAP-induced acute liver injury timecourse; 3) ST spots human liver (n=3 healthy, n=2 APAP-ALF, n=2 NAE-ALF) and mouse liver (n=1 per time point) multiplex smFISH (n=2 healthy human liver and n=2 APAP-ALF liver samples).

### snRNA-seq analysis

We aligned to GRCh38 and mm10 (Ensembl 93) reference genomes (modified to allow intronic as well as exonic feature alignment), and estimated nuclei-containing partitions and unique molecular identifiers (UMIs), using the CellRanger v3.1.0 Single-Cell Software Suite from 10X Genomics.

To enable reproducible analysis, we developed the SeuratPipe R package v1.0.0 (https://doi.org/10.5281/zenodo.7331092), a pipeline building on existing packages. In brief we performed analysis as follows:

We performed per-dataset quality control in the Seurat R package v4.1.1. We used the Scrublet python module v0.2.3^38^ to identify potential doublets and the SoupX^39^ R package v1.5.2 to automatically calculate and correct for background contamination. Finally, we excluded nuclei that expressed fewer than 1000 genes, or mitochondrial gene content >5% of the total UMI count.

After merging the individual datasets, we normalised feature counts per nuclei by dividing the total UMI count for that nuclei, then multiplying by a scale factor of 10000 and natural-log transforming. We corrected for sample bias by obtaining PC embeddings using the Harmony R package v0.1.0^40^. Further, we downsampled the hepatocyte populations to standardise sample contribution to downstream analysis.

Nuclei clusters were identified using the shared nearest neighbour (SNN) modularity optimization-based clustering algorithm implemented in *Seurat*, determining dataset variability for the purpose of constructing the SNN graph using Harmony-corrected principal components. We calculated differentially-expressed features using a Wilcoxon Rank Sum test. To annotate these clusters, we used a curated list of known marker genes per cell lineage in the liver (SI Table 2) to obtain signature scores using the *AddModuleScore* function in *Seurat*. Clusters identified as primarily composed of cycling cells were re-clustered to split them out into their constituent lineages as above. We then iteratively applied the above workflow for each lineage thus identified, inserting a ‘cleansing’ step where we removed clusters displaying an abundance of nuclei previously identified as doublets or overexpressing marker genes of other lineages. We generated a hepatocyte migration gene module (SI Table 2) using the top 25 (by avg_log2FC) differentially-expressed features from the human migratory hepatocyte cluster. Gene ontology analysis was performed using the *clusterProfiler* R package^41^. Liver zonation specificity scores were obtained by first scaling central and portal zonation signature scores to a value between 0 and 1, and subsequently setting *zonation_score = central_score / (central_score + portal_score)*.

### Spatial transcriptomics analysis

We aligned to GRCh38 and mm10 (Ensembl 93) reference genomes using the SpaceRanger v1.0.0 Spatial Gene Expression Software Suite from 10X Genomics.

We performed per-dataset quality control in the *Seurat* R package v4.1.1. We excluded spots expressing fewer than 800 genes, or mitochondrial gene content >20% for both human and mouse samples of the total UMI count. We also manually filtered low quality spots (those isolated from main tissue section) using the 10X Genomics Loupe browser (v5.0). Similar to snRNA-seq we computed gene signature scores for hepatocytes (*TTR, TF, HP, CYP2A6, CYP2E1, CYP3A4, HAL*), myofibroblasts (*ACTA2, COL1A1, COL1A2, COL3A1*), and cycling cells (genes listed in *Seurat* cc.genes.updated.2019).

Gene expression and tissue topography were used to draw spatial trajectories across healthy and APAP-ALF tissues via the *SPATA2* R package v0.1.0^42^. The trajectory modelling functionality of *SPATA2* was used to identify central- and portal-associated genes and corresponding modules (SI Table 2) whose expression trajectory followed the underlying spatial model.

### Multiplex smFISH analysis

Nuclei segmentation and expansion was performed using QuPath to demarcate cells. A gene-cell matrix was then obtained and signature scores in tissue were computed using pre-defined cell populations.

### Further statistical analysis

Further statistical analyses were performed using GraphPad Prism. Comparison of changes between two groups was performed using a two-tailed paired t-test or unpaired t-test. Comparison of changes between groups was performed using a two-way ANOVA with Sidaks multiple comparison test with a single pooled variance. *P* < 0.05 was considered statistically significant.

## Data availability

Our snRNA-seq and spatial transcriptomics data are freely available for user-friendly interactive browsing online at https://shiny.igc.ed.ac.uk/7efa5350ba94425388e47ea7cdd5aa64/.

All raw and processed sequencing data are deposited in the Gene Expression Omnibus (GEO) under accession number GSE223561.

## Code availability

All code will be made available upon acceptance of the manuscript by the Human Cell Atlas.

## Acknowledgements

This work was supported by a Wellcome Trust Senior Research Fellowship in Clinical Science (ref. 219542/Z/19/Z) to N.C.H., a Chan Zuckerberg Initiative Seed Network Grant to N.C.H., and a Tenovus Scotland grant (E20-03) to K.P.M., S.J.W. and N.C.H. J.P. was supported by a Medical Research Council Precision Medicine PhD studentship. C.A.K. is a cross-disciplinary post-doctoral fellow (XDF) supported by funding from the University of Edinburgh and Medical Research Council (MC_UU_00009/2). T.G.B. was funded by the Wellcome Trust (ref. WT107492Z). S.M. was funded by Cancer Research UK core funding to the CRUK Beatson Institute (ref. A17196 and A31287). F.F., J.B.G.M and L.M.C. were supported by CRUK core funding to the Beatson Institute A31287 and CRUK core funding to L.M.C. A23983. P.R. was supported by an MRC Clinician Scientist Fellowship (MR/N008340/1) and MRC Senior Clinical Fellowship (MR/W015919/1). We thank the patients who donated liver tissue for this study. We thank J. Davidson, J. Black, C. Ibbotson and A. Baird of the Scottish Liver Transplant Unit and the research nurses of the Wellcome Trust Clinical Research Facility for assistance with consenting patients for this study. We thank the liver transplant coordinators and surgeons of the Scottish Liver Transplant Unit and the surgeons and staff of the Hepatobiliary Surgical Unit, Royal Infirmary of Edinburgh for assistance in procuring human liver samples. We thank core facilities and services at the Beatson Institute, in particular the Biological Research Unit & the Beatson Advanced Imaging Resource (BAIR). We thank Guillaume Jacquemet (Cell Migration Lab, Turku, Finland) for advice on Cellpose segmentation for IVM analysis. We thank Robert Insall and Laura Machesky for helpful scientific discussions. We thank W. Mungall for technical support. We thank C. Nicol for help with figure illustrations. We thank N. Pham for technical assistance with Incucyte. We thank Catherine Winchester and Ruoyan Li for critical reading of the manuscript. We gratefully acknowledge the contribution to this study made by the University of Birmingham’s Human Biomaterials Resource Centre which has been supported through Birmingham Science City -Experimental Medicine Network of Excellence project.

This research was funded in whole, or in part, by the Wellcome Trust (Wellcome Trust Senior Research Fellowship in Clinical Science to N.C.H. ref. 219542/Z/19/Z). For the purpose of open access, the author has applied a CC BY public copyright licence to any Author Accepted Manuscript version arising from this submission.

## Author contributions

K.P.M. performed experimental design, data generation and data analysis and interpretation; J.R.W-K., J.R.P., and C.A.K. performed computational analyses on the human snRNA-seq data: J.R.P. analysed hepatocytes, J.R.W-K. analysed non-hepatocyte lineages. J.R.W-K., J.R.P., and C.A.K. performed computational analyses on the mouse snRNA-seq data: J.R.P. analysed hepatocytes, J.R.W-K. analysed non-hepatocyte lineages. C.A.K. performed computational analyses on the spatial transcriptomics data. C.A.K. and K.P.M. performed computational analysis on the multiplex smFISH data. C.A.K. developed the *SeuratPipe* package and generated the interactive online browser. K.P.M., F.F., J.B.G.M., S.M., T.G.B., N.C.H. and L.M.C. performed experimental design, data generation, data analysis and interpretation for the mouse liver intravital microscopy experiments. M.B., E.Z., M.B-S., E.S., G.W., N.T.Y. and R.D. performed data generation and analysis; S.J.W., L.K., G.C.O., S.J.W. and P.R. procured human liver tissue; N.O.C. provided support for the cell migration analyses; K.J.S. procured human liver tissue and provided intellectual contribution; T.J.K. provided liver pathology expertise; the Acute Liver Failure Study Group, J.R., W.M.L., M.H., and C.J.W. provided ALF human liver tissue; C.A.V., J.C.M. and S.A.T provided advice with computational analyses; T.G.B. provided support with hepatocyte lineage tracing, mouse liver intravital microscopy experiments and data interpretation; L.C. provided expertise with mouse liver intravital microscopy experiments and performed data interpretation; K.P.M., J.R.W-K., J.R.P., C.A.K. and N.C.H. wrote the manuscript. N.C.H. conceived the study, designed experiments, interpreted data and supervised the study.

## Competing interest declaration

N.C.H. has received research funding from AbbVie, Pfizer, Gilead, Boehringer-Ingelheim and Galecto, and is an advisor or consultant for Astra-Zeneca, Galecto, GSK, MSD, Pliant Therapeutics, Ambys Medicines, Mediar Therapeutics and Q32 Bio.

Supplementary Information is available for this paper.

Correspondence and requests for materials should be addressed to Neil Henderson.

## Extended data figure legends

**Extended Data Figure 1:**
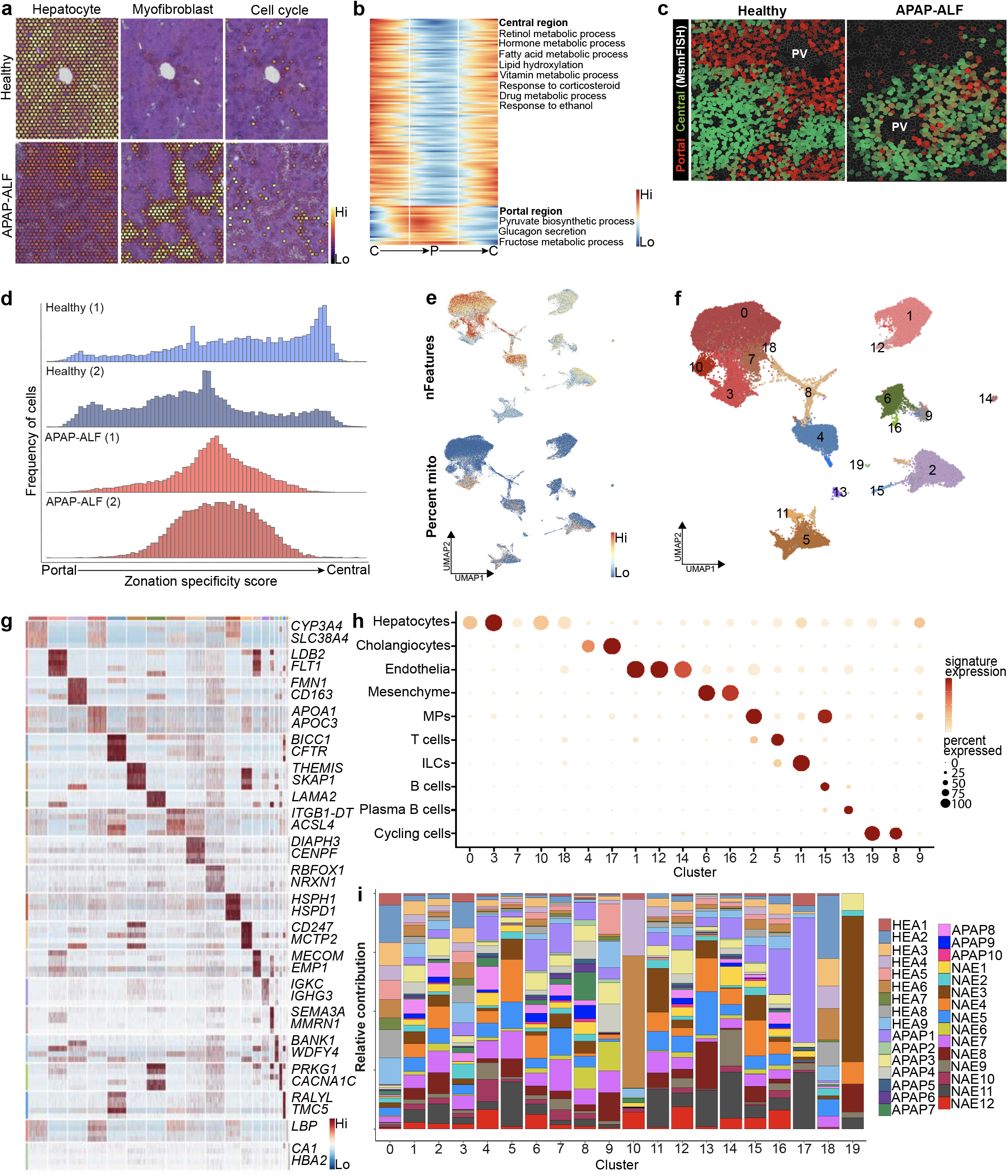
Disruption of zonation and lineage annotation in human acute liver failure. **a**, Spatial expression (ST) of hepatocyte, myofibroblast, and cell cycle signatures across healthy and APAP-ALF liver. **b**, Differential GO terms across spatial trajectory analysis from peri-central to peri-portal to peri-central regions in healthy human liver, with exemplar terms labelled (right). **c**, Spatial expression (multiplex smFISH) of known human zonal gene modules in healthy (n=2) and APAP-ALF (n=2) liver tissue. **d**, Distribution of zonal specificity score in healthy (n=2) and APAP-ALF (n=2) human liver tissue. **e**, Quality control metrics (mitochondrial percentage, number of features) across the human snRNA-seq dataset. **f**, UMAP visualisation of 72,262 nuclei from healthy (n=9), APAP-ALF (n=10), and NAE-ALF (n=12) human liver explants, annotated by clustering. **g**, Heatmap of marker genes (colour-coded by cluster) with exemplar genes labelled (right). Columns denote cells, rows denote genes. **h**, Dotplot annotating clusters by lineage signature expression. Circle size indicates cell fraction expressing signature greater than mean; colour indicates mean signature expression. **i**, Stacked bar plot denoting relative contribution of liver sample to each cluster.

**Extended Data Figure 2:**
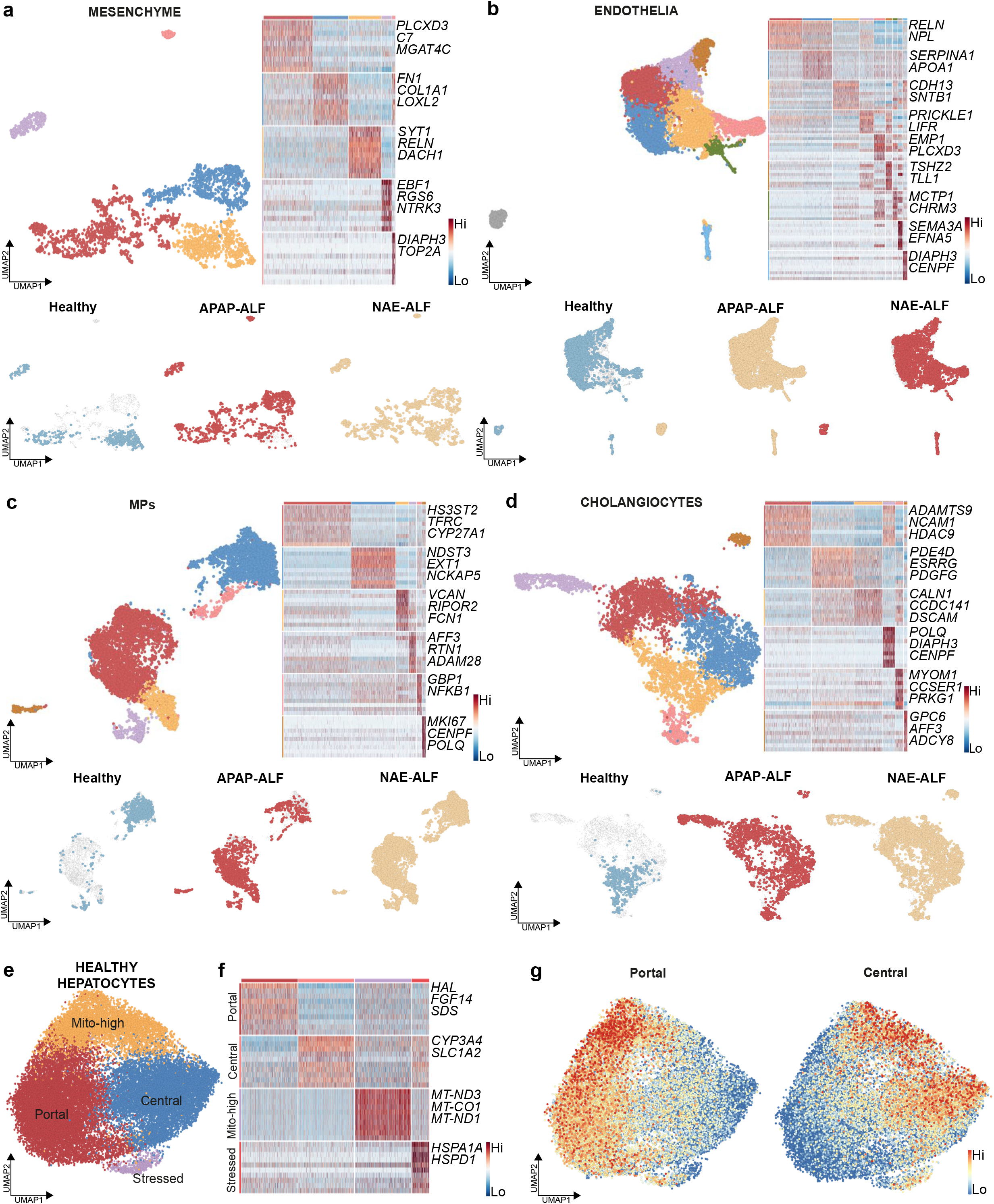
Lineage analysis of human liver regeneration atlas. UMAPs of (**a**) mesenchyme, (**b**) endothelia, (**c**) MPs and (**d**) cholangiocytes derived from human liver single nuclei RNA sequencing, coloured by clustering (top left) and aetiology (bottom); heatmap (right) of marker genes (colour-coded by cluster) with exemplar genes labelled (right), columns denote cells, rows denote genes. **e**, UMAP of healthy human hepatocyte nuclei, coloured by cluster. **f**, Heatmap of marker genes in healthy hepatocyte clusters (colour-coded by cluster) with exemplar genes labelled (right). Columns denote cells, rows denote genes. **g**, Human SPATA-derived portal and central region gene modules applied to healthy hepatocytes.

**Extended Data Figure 3:**
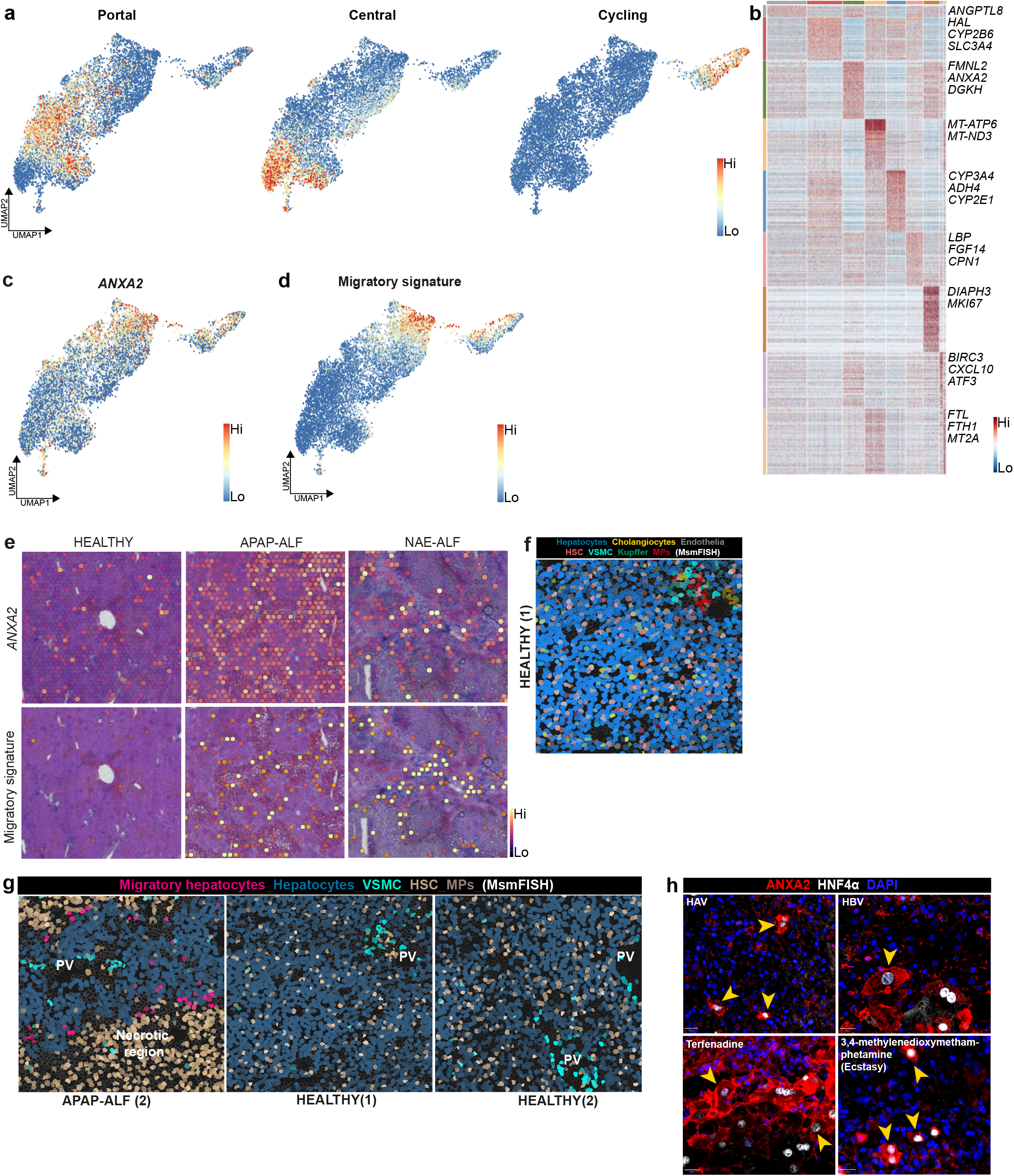
Migratory hepatocytes in human acute liver failure. **a**, UMAP of human hepatocyte nuclei, all aetiologies, showing portal, central and cycling gene module scores. Gene modules in SI Table 2. **b**, Heatmap of marker genes in hepatocyte clusters (colour-coded by cluster) with exemplar genes labelled (right). Columns denote cells, rows denote genes. **c**, UMAP of human hepatocyte nuclei, all aetiologies, showing *ANXA2* gene expression. **d**, UMAP of human hepatocyte nuclei, all aetiologies, showing migratory gene module signature (SI Table 2). **e**, Spatial expression of migratory signature (gene modules in SI Table 2) and ANXA2 in healthy, APAP-ALF and NAE-ALF human liver tissue. **f**, Multiplex smFISH showing cell lineages in healthy human liver tissue. Gene modules in SI Table 2. **g**, Multiplex smFISH showing migratory hepatocytes in APAP-ALF human liver in relation to other cell lineages. Gene modules in SI Table 2. **h**, Representative immunofluorescence images of ANXA2 (red), HNF4α (hepatocytes, white) and DAPI (nuclear stain, blue) in hepatitis A-induced ALF, hepatitis B-induced ALF, and drug-induced ALF. Yellow arrowheads denote ANXA2+ hepatocytes with migratory phenotype. Scale bar 20µm.

**Extended Data Figure 4:**
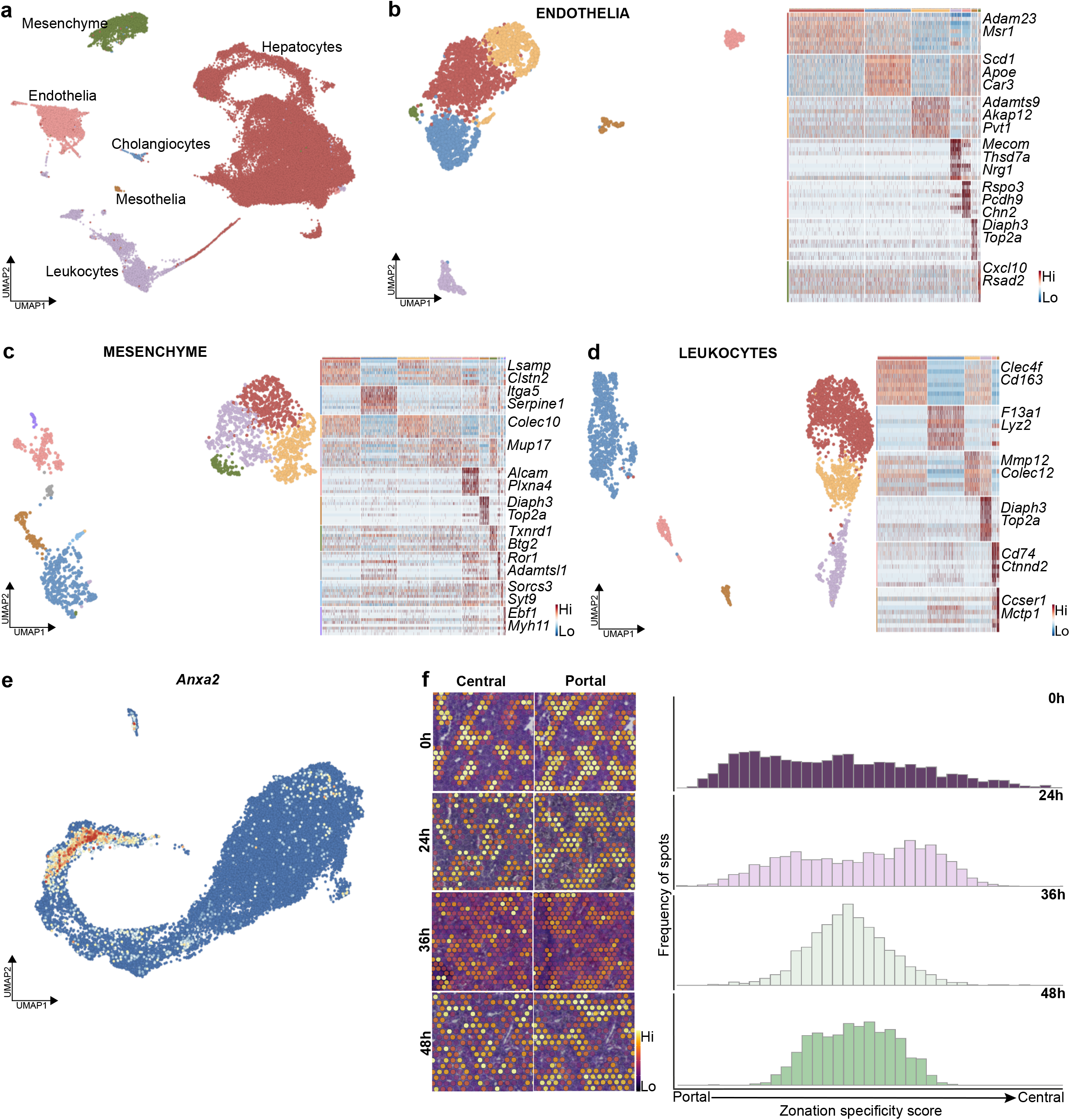
Mouse APAP-induced acute liver injury. **a**, UMAP visualisation of mouse liver nuclei, annotated by lineage inferred using signatures of known lineage markers (SI Table 2). **b**, UMAP (left) of mouse endothelia annotated by cluster. Heatmap (right) of marker genes in clusters (colour-coded by cluster) with exemplar genes labelled. Columns denote cells, rows denote genes. **c**, UMAP (left) of mouse mesenchyme annotated by cluster. Heatmap (right) of marker genes in clusters (colour-coded by cluster) with exemplar genes labelled. Columns denote cells, rows denote genes. **d**, UMAP (left) of mouse leukocytes annotated by cluster. Heatmap (right) of marker genes in clusters (colour-coded by cluster) with exemplar genes labelled. Columns denote cells, rows denote genes. **e**, UMAP of mouse hepatocyte nuclei, all time points, showing *Anxa2* gene expression. **f**, Spatial expression (ST) in selected timepoints post APAP-induced mouse liver injury (left) of mouse liver-derived zonal gene modules. Distribution of zonal scores across selected timepoints post APAP-induced mouse liver injury (right).

**Extended Data Figure 5:**
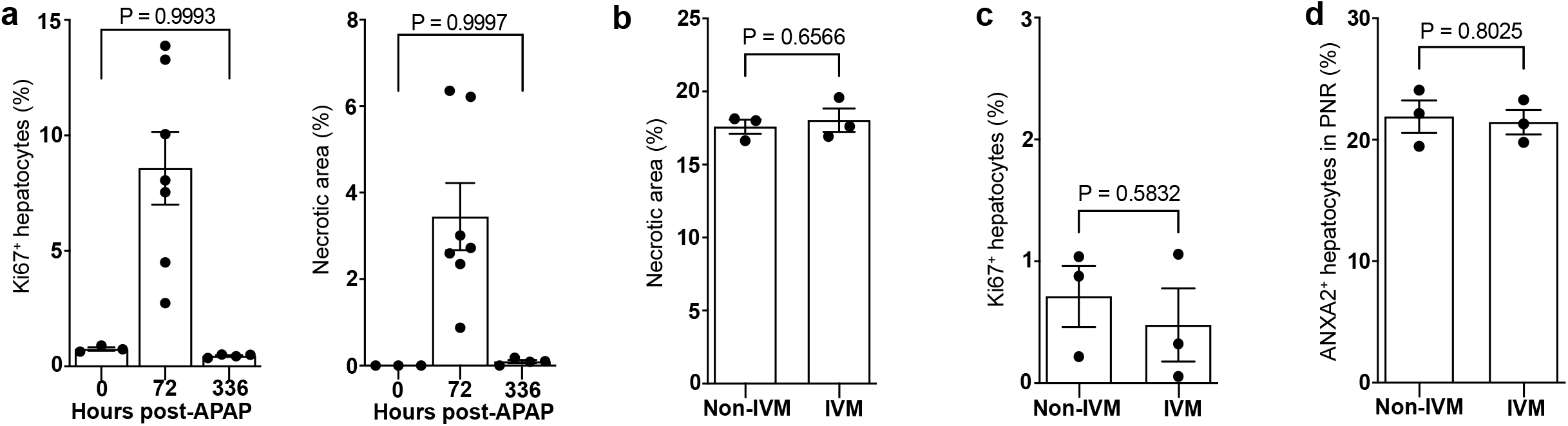
APAP-induced acute liver injury is not altered by intravital microscopy (IVM) **a**, Quantification of hepatocyte proliferation (left) and necrotic area (right) following APAP-induced mouse liver injury. Unpaired t-test, n=3 (0h), n=7 (72h), n=4 (336h). Data are mean±SEM. Quantification of necrotic area (**b**), Ki67+ hepatocytes (**c**), and ANXA2+ hepatocytes in the PNR (**d**), following APAP-induced mouse liver injury in IVM and non-IVM (42 hours) mice. Unpaired t-test, n=3. Data are mean±SEM.

**Extended Data Figure 6:**
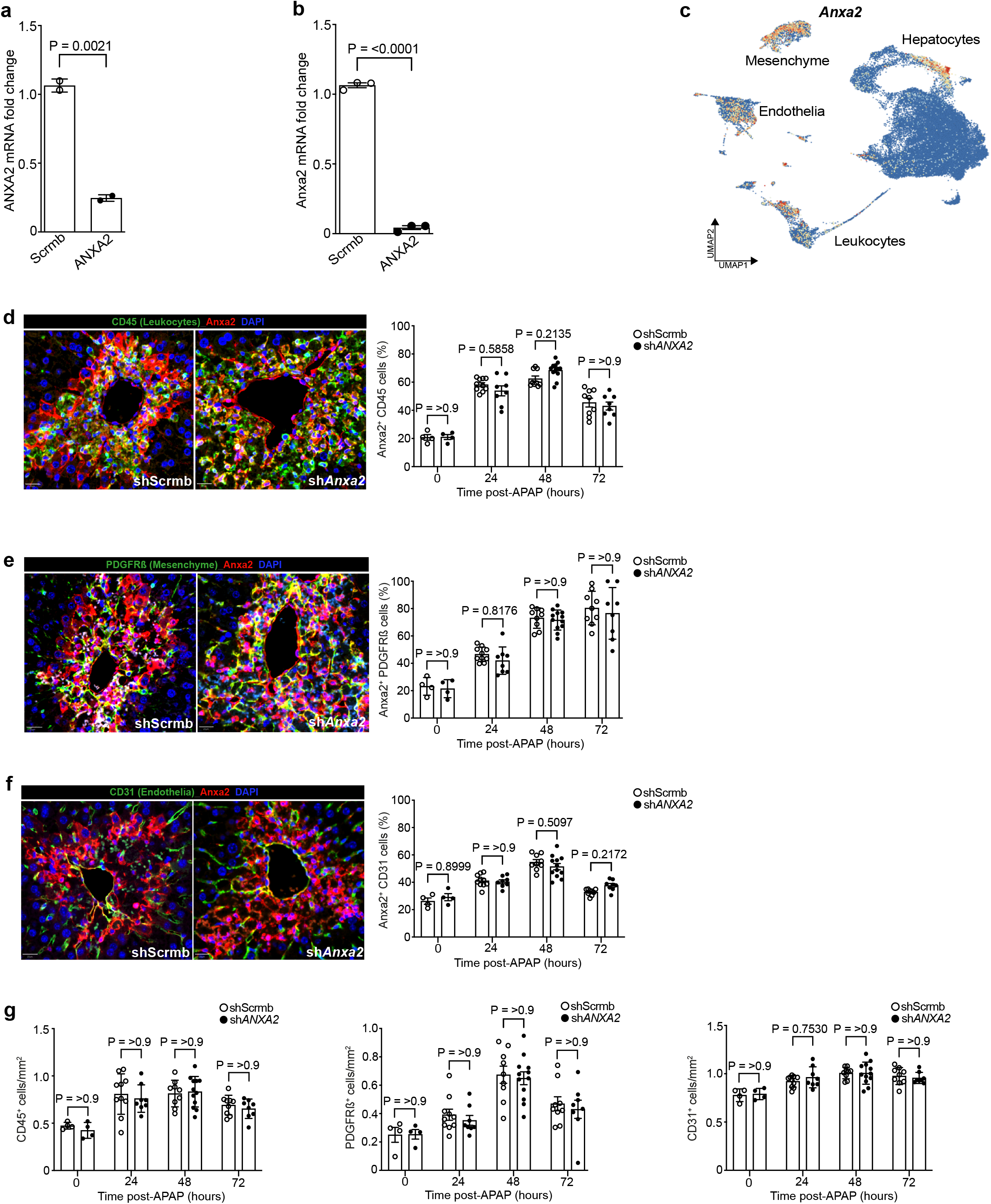
AAV8-shRNA-ANXA2 does not affect Anxa2 expression in non-hepatocyte lineages. **a**, *ANXA2* gene expression (RT-qPCR analysis) in Huh7 cells (human hepatocyte cell line) treated with Scrmb (control) or *ANXA2* siRNA. Unpaired t-test, mean±SEM, two independent experiments. **b**, *Anxa2* gene expression (RT-qPCR) in mouse primary hepatocytes treated with Scrmb (control) or *Anxa2* siRNA. Unpaired t-test, mean±SEM, three independent experiments. **c**, UMAP visualisation of *Anxa2* gene expression in mouse APAP-injured liver nuclei. **d**, Representative immunofluorescence images of CD45 (leukocytes, green), ANXA2 (red) and DAPI (nuclear stain, blue) in AAV8-shRNA-Scrmb or AAV8-shRNA-*Anxa*2 treated mice 48hrs post APAP-induced acute liver injury (left). Scale bar 20µm. Quantification of CD45^+^/ANXA2^+^ cells following APAP-induced acute liver injury in AAV8-shScrmb or AAV8-sh*Anxa*2 treated mice (right). Two-way ANOVA, n=4 (0h), n=8 (24h), n=11 (48h), n=8 (72h). Data are mean±SEM. **e**, Representative immunofluorescence staining of PDGFRβ (mesenchyme, green), ANXA2 (red) and DAPI (nuclear stain, blue) in AAV8-shRNA-Scrmb or AAV8-shRNA-*Anxa*2 treated mice 48hrs post APAP-induced acute liver injury (left). Scale bar 20µm. Quantification of PDGFRβ^+^/ANXA2^+^ cells following APAP-induced acute liver injury in AAV8-shScrmb or AAV8-sh*Anxa*2 treated mice (right). Two-way ANOVA, n=4 (0h), n=8 (24h), n=11 (48h), n=8 (72h). Data are mean±SEM. **f**, Representative immunofluorescence staining of CD31 (endothelia, green), ANXA2 (red) and DAPI (nuclear stain, blue) in AAV8-shRNA-Scrmb or AAV8-shRNA-*Anxa*2 treated mice 48hrs post APAP-induced acute liver injury (left). Scale bar 20µm. Quantification of CD31^+^/ANXA2^+^ cells following APAP-induced acute liver injury in AAV8-shScrmb or AAV8-sh*Anxa*2 treated mice (right). Two-way ANOVA, n=4 (0h), n=8 (24h), n=11 (48h), n=8 (72h). Data are mean±SEM. **g**, Quantification of PDGFRβ^+^, CD45^+^ or CD31^+^ cells/mm^2^ following APAP-induced acute liver injury in AAV8-shScrmb or AAV8-sh*Anxa*2 treated mice. Two-way ANOVA, n=4 (0h), n=8 (24h), n=11 (48h), n=8 (72h). Data are mean±SEM.

## Supplementary information legends

**Supplementary Information Video 1:**

Application of human migratory hepatocyte gene module to mouse hepatocyte nuclei across timepoints post APAP-induced acute liver injury.

**Supplementary Information Video 2:**

Application of *Anxa2* gene expression to mouse hepatocyte nuclei across timepoints post APAP-induced acute liver injury.

**Supplementary Information Video 3:**

Application of SPATA-derived mouse central (left) and portal (right) zonal signatures to mouse hepatocyte nuclei across timepoints post APAP-induced mouse liver injury.

**Supplementary Information Video 4:**

Intravital imaging of mouse liver in *Hep;tdTom* reporter mice (hepatocytes express cytoplasmic tdTomato) demonstrating centrilobular hepatocyte necrosis in real-time from 24 hours post APAP-induced mouse liver injury. Scale bar 30µm.

**Supplementary Information Video 5:**

Intravital imaging of mouse liver in *Hep;tdTom* reporter mice (hepatocytes express cytoplasmic tdTomato) demonstrating centrilobular hepatocyte necrosis in real-time from 24 hours post APAP-induced mouse liver injury. Scale bar 30µm.

**Supplementary Information Video 6a:**

Intravital imaging of mouse liver in *Hep;tdTom* reporter mice (hepatocytes express cytoplasmic tdTomato) demonstrating wound closure between 36-42 hours following APAP-induced mouse liver injury. Boxed area is magnified in SI Video 6b. Scale bar 30µm.

**Supplementary Information Video 6b:**

Magnification of boxed area from SI Video 6a -intravital imaging of mouse liver in *Hep;tdTom* reporter mice (hepatocytes express cytoplasmic tdTomato) demonstrating wound closure between 36-42 hours following APAP-induced mouse liver injury. Scale bar 30µm.

**Supplementary Information Video 7:**

Intravital imaging of mouse liver in *Hep;tdTom* reporter mice (hepatocytes express cytoplasmic tdTomato) between 36-42 hours following APAP-induced mouse liver injury. White arrowheads denote hepatocytes with a motile morphology, including membrane ruffling and the formation of lamellipodia at the hepatocyte leading edge abutting the wound. Scale bar 5µm.

**Supplementary Information Video 8:**

Intravital imaging of mouse liver in *Hep;tdTom* reporter mice (hepatocytes express cytoplasmic tdTomato) between 36-42 hours following APAP-induced mouse liver injury. White arrowheads denote hepatocytes with a motile morphology, including membrane ruffling and the formation of lamellipodia at the hepatocyte leading edge abutting the wound. Scale bar 10µm.

**Supplementary Information Video 9:**

Intravital imaging of mouse liver in *Hep;tdTom* reporter mice (hepatocytes express cytoplasmic tdTomato) between 36-42 hours following APAP-induced mouse liver injury. White arrowheads denote hepatocytes with a motile morphology, including membrane ruffling and the formation of lamellipodia at the hepatocyte leading edge abutting the wound. Scale bar 5µm.

**Supplementary Information Video 10:**

Intravital imaging of mouse liver in *Hep;tdTom* reporter mice (hepatocytes express cytoplasmic tdTomato) between 36-42 hours following APAP-induced mouse liver injury. White arrowheads denote hepatocytes with a motile morphology, including membrane ruffling and the formation of lamellipodia at the hepatocyte leading edge abutting the wound. Scale bar 5µm.

**Supplementary Information Video 11:**

Intravital imaging of mouse liver in *Hep;tdTom* reporter mice (hepatocytes express cytoplasmic tdTomato) between 36-42 hours following APAP-induced mouse liver injury. White arrowheads denote hepatocytes with a motile morphology, including membrane ruffling and the formation of lamellipodia at the hepatocyte leading edge abutting the wound. Scale bar 5µm.

**Supplementary Information Video 12:**

Intravital imaging of mouse liver in *Hep;tdTom* reporter mice (hepatocytes express cytoplasmic tdTomato) between 36-42 hours following APAP-induced mouse liver injury. White arrowheads denote hepatocytes with a motile morphology, including membrane ruffling and the formation of lamellipodia at the hepatocyte leading edge abutting the wound. Scale bar 5µm.

**Supplementary Information Video 13:**

Intravital imaging of mouse liver in *Hep;tdTom* reporter mice (hepatocytes express cytoplasmic tdTomato) between 36-42 hours following APAP-induced mouse liver injury. White arrowheads denote hepatocytes with a motile morphology, including membrane ruffling and the formation of lamellipodia at the hepatocyte leading edge abutting the wound. Scale bar 5µm.

**Supplementary Information Table 1:**

Patient metadata (snRNAseq and Spatial Transcriptomics).

**Supplementary Information Table 2:**

Gene signatures in human and mouse snRNA-seq and ST, and human multiplex smFISH.

**Supplementary Information Table 3:**

Marker genes for human and mouse snRNA-seq.

**Supplementary Information Table 4:**

GO results for human and mouse snRNA-seq, and ST SPATA analysis.

**Supplementary Information Table 5:**

Antibodies and reagents.

